# Mammalian CST averts replication failure by preventing G-quadruplex accumulation

**DOI:** 10.1101/354050

**Authors:** Yang Liu, Miaomiao Zhang, Bing Wang, Yingnan Xiao, Tingfang Li, X Geng, Guang Li, Qiang Liu, Carolyn M. Price, Feng Wang

## Abstract

Human CST (CTC1-STN1-TEN1) is an RPA-like complex that associates with G-rich single-strand DNA and helps resolve replication problems both at telomeres and genome-wide. We previously showed that CST binds and disrupts G-quadruplex (G4) DNA *in vitro*, suggesting that CST may prevent *in vivo* blocks to replication by resolving G4 structures. Here, we demonstrate that CST binds and unfolds G4 with similar efficiency to RPA. In cells, CST is recruited to telomeric and non-telomeric chromatin upon G4 stabilization. STN1 depletion increases G4 accumulation and slows bulk genomic DNA replication. At telomeres, combined STN1 depletion and G4 stabilization causes multi-telomere FISH signals and telomere loss, hallmarks of deficient telomere duplex replication. Strand-specific telomere FISH indicates preferential loss of C-strand DNA while analysis of BrdU uptake during leading and lagging-strand telomere replication shows preferential under-replication of lagging telomeres. Together these results indicate a block to Okazaki fragment synthesis. Overall, our findings indicate a novel role for CST in maintaining genome integrity through resolution of G4 structures both ahead of the replication fork and on the lagging strand template.

## Introduction

Telomeres are specialized nucleoprotein structures that maintain genome stability by preventing chromosome ends from being detected as DNA damage and triggering end-to-end fusion through unwanted DNA repair reactions[1–3]. In mammalian cells, the telomeric DNA consists of thousands of TTAGGG•AATCC repeats which terminate with a short region of ssDNA on the G-rich strand. This repeated sequence DNA is packaged by shelterin, a six protein complex that sequesters the DNA terminus to prevent inappropriate DNA damage signaling[4, 5]. Mammalian telomeres also associate with a ssDNA-binding complex called CST (CTC1-STN1-TEN1) that is essential for multiple aspects of telomere replication[6–8]. Telomeres are particularly challenging to replicate due their G-rich sequence and terminal structure[3, 9]. The repetitive G-rich DNA tends to stall the replication fork, while the inability of DNA polymerase to replicate the DNA 5’ end leads to progressive telomere shortening unless additional sequence is added to the DNA terminus. This telomere elongation is achieved by telomerase which extends the G-strand and DNA polymerase which then synthesizes the complementary C-strand.

The CTC1 and STN1 subunits of CST were originally identified as DNA polymerase α-primase (Pol α) cofactors that increase the affinity of Pol α for ssDNA templates[10]. More recent studies have shown that one role of CST is in regulation of Pol α switching from RNA and DNA synthesis[11], a process that is essential to generate Okazaki fragments during lagging strand replication. *In vivo* studies have shown that CST participates in multiple stages of telomere replication. First, it aids in replication of the G-rich DNA duplex. Depletion of CST subunits slows replication through this region and leads to telomere fragility and sudden telomere loss, suggesting that CST either prevents replication fork stalling or facilitates fork restart[12–14]. CST then regulates telomerase to prevent G-strand overextension([15–17]). Finally, CST is required to engage Pol y for C-strand synthesis so that the ssDNA generated by telomerase is converted to new telomere duplex[12, 16, 18, 19]. In the absence of CST, the lack of C-strand synthesis leads to progressive telomere shortening with each round of replication, similar to a telomerase knockout. CST also has less well understood non-telomeric roles related to recovery from replication stress[20, 21]. When hydroxylurea is used to stall replication, CST promotes replication restart by increasing the firing of dormant replication origins[14, 21]. More recently, CST was shown to localize to GC-rich loci in response to replication stress where it functions to prevent fragile site expression[22].

CST is structurally similar to Replication Protein A (RPA), the ssDNA binding protein that directs DNA replication, repair, and recombination in eukaryotic cells[23–25]. The structural conservation encompasses the OB-folds, winged helix domains and the dimerization interface of the two smaller subunits (STN1-TEN1 and RPA2-RPA3). Like RPA, CST appears to contact DNA via multiple OB folds leading to a dynamic mode of DNA binding. Binding and release of individual OB fold is thought to underlie the capacity of both proteins to melt DNA structure and regulate DNA association and activity of interaction partners such as DNA polymerases. Although CST and RPA share many similarities in terms of structure and DNA binding activity, they are architecturally different and they play quite distinct roles during DNA replication and repair[26]. Moreover, CST binds preferentially to G-rich ssDNA and, unlike ssDNA-bound RPA, it does not trigger ATR activation[19, 25, 27]. Thus, CST appears to have evolved a unique role by serving to resolve a variety of replication problems without activation of DNA damage signaling.

Currently the mechanism by which CST facilitates replication through G-rich dsDNA is unclear. However, the recent discovery that CST can unwind G-quadruplex (G4) DNA[26], suggests that the role of CST is to prevent or resolve replication fork stalling at G4 structures. G4s form in G-rich DNA due to the ability of guanine to form thermodynamically stable Hoogsteen base-paired quartets[28]. The human genome contains thousands of regions with G4-forming potential[29]. These regions are becoming appreciated as an important genomic feature because of their capacity to form stable G4 structures that interrupt the DNA duplex[30, 31]. The G4 DNA is thought to play roles in various cellular processes including transcriptional activation, stimulation of meiotic recombination, chromatin assembly and rudimentary telomere capping[32, 33]. Despite serving these beneficial regulatory functions, G4 structures can also have adverse effects on genome stability because they impose a structural barrier to DNA replication. Any G4 structures formed as a result of transcription may later block passage of the DNA replication machinery[34, 35]. Replication may also be blocked by G4 formed when ssDNA is exposed during lagging strand synthesis.

The recent discovery that CST can resolve G4 DNA *in vitro*, suggested that CST may also remove G4 structures *in vivo* and this property could explain the ability of CST to facilitate replication through telomeres and other G-rich regions. We now provide evidence for this by showing that CST is needed to prevent G4 accumulation and STN1/CST loss slows bulk DNA replication after G4 stabilization. At telomeres, combined STN1 depletion and G4 stabilization causes replication-associated defects on both sister chromatids but lagging-strand telomere replication is most heavily affected. Our results indicate that the G4 resolving activity of CST is necessary to prevent a range of replication defects.

## Results

### CST and RPA unfold telomeric G-quadruplex DNA with similar efficiency

Although both CST and RPA can unfold G4 DNA[26, 36], their relative efficiency in removal of this structure has not been examined. Thus, as a first step towards understanding the biological importance of the CST G4 resolving activity, we directly compared the ability of CST and RPA to bind and unfold G4 DNA formed by a telomeric oligonucleotide Tel21 (GGG(TTAGGG)_3_. In initial experiments we used circular dichroism to confirm Tel21 folding into a G4 structure (Fig. S1). Consistent with previous reports[37], when Tel21 was incubated in buffer containing 150 mM NaCl, the CD revealed a trough at 265 nm and peak at 295 nm indicating G4 formation. As expected, incubation in 150 mM LiCl destabilized the G4 structure. We next used electrophoretic mobility shift assays (EMSA) to compare CST and RPA binding to Tel21 in the presence of either 150 mM LiCl or NaCl (Fig. 1A-E). ^32^P-labeled Tel21 was incubated in LiCl or NaCl containing buffer for 30 min to allow DNA folding/unfolding, purified CST or RPA was added for 30 min, then samples were separated in native agarose gels. The EMSA analysis indicated that in conditions favoring ssDNA over G4 formation (150 M LiCl), CST and RPA bound Tel21 with similar efficiency (Kd(app) 1.3 nM for CST versus 1.9 nM for RPA). However, in 150 nM NaCl, where Tel21 formed a G4, CST showed slightly lower affinity than RPA (Kd(app) 8.5 nM for CST versus 5.5 nM for RPA). (Fig. 1B, D & E).)

**Figure 1.**
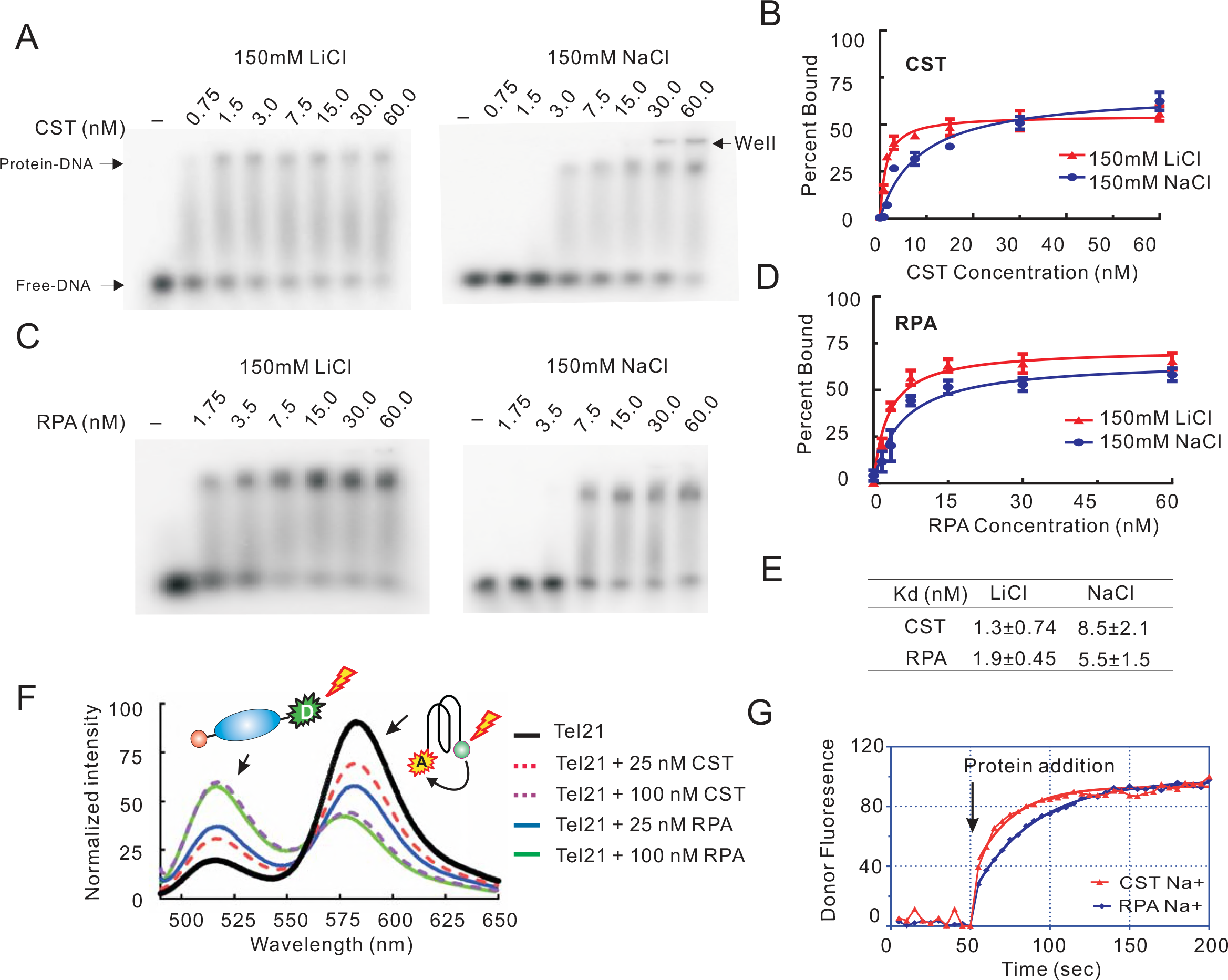
CST and RPA bind telomeric ssDNA and disrupt G4 structure. (A-E) Comparison of CST and RPA binding to Tel21 in 150 mM LiCl or NaCl. (A & C) EMSAs showing CST (A) or RPA (C) binding. Reactions contained the indicated amounts of protein and 0.05 nM DNA. (B & D) Quantification of CST (B) or RPA (D) binding to Tel21 in LiCl versus NaCl. Data were fit to a one site specific model to obtain binding isotherms and Kd(app). Mean ± SEM, n = 3 independent experiments each with a different protein preparation. (E) Comparison of Kd(app) for CST and RPA binding to Tel21 in LiCl or NaCl. (F) Emission spectra for 5’(FAM)-Tel21-(TAMRA)3’ following addition of indicated amount of CST or RPA and excitation with 494 nM light. Reactions contained 150 mM KCl. (F) FRET showing the disruption of G4 by CST and RPA. Donor fluorescence monitored in real-time upon addition of 100 nM CST or RPA. The base line was set to 0 and the maximum value for scan was set to 100. CST or RPA was added at the 50 seconds time point. Related to Figure S1.

Since CST bound G4 DNA, we next used Fluorescence Resonance Energy Transfer (FRET) to compare the capacity of CST and RPA to unfold G4 structure. The FRET substrate was double-labeled Tel21 that had fluorescein (FAM) at the 5’ end as the FRET donor and tetramethylrhodamine (TAMRA) at the 3’ end as the FRET receptor (Fig. 1F top). As we previous demonstrated, when this oligomer forms a G4, the two fluorophores become closely juxtaposed, leading to FRET and suppression of the donor fluorescence[38]. To assess G4 unfolding, we added either CST or RPA to the folded Tel21, excited the donor fluorophore with 480 nm light and examined the emission spectrum. As anticipated, addition of RPA caused a concentration-dependent reduction in FRET, as seen by an increase in donor emission (518 nm) and a concomitant decrease in acceptor emission (580 nm). This loss of FRET indicated increased distance between the two fluorophores due to G4 unfolding[38]. Addition of CST also resulted in a concentration-dependent decrease in donor emission and an increase in acceptor emission indicating that, like RPA, CST can unfold G4 structure.

To compare the kinetics of G4 unfolding by CST and RPA, we performed a time course study where donor fluorescence was measured before and after CST or RPA addition. Double-labeled Tel21 was incubated in 150 mM NaCl to promote G4 formation, then 100 nM protein was added to maximize unfolding. To our surprise, we found that CST caused a faster increase in donor fluorescence compared to RPA (Fig. 1F). Thus, although CST has a slightly lower affinity for Tel21 G4 DNA than RPA, CST appears to initiate G4 opening more rapidly at saturating protein concentrations. Despite the subtle differences in Kd(app) and rate of G4 unfolding, our data indicate that CST and RPA generally recognize and unfold G4 DNA with similar efficiency. This finding suggests that CST and RPA are equally well equipped for *in vivo* G4 removal. Although, CST is less abundant than RPA, local concentrations may be quite high as CST interacts with the telomere protein TPP1 and itis enriched at telomeres and other G-rich regions[15, 39].

### CST is recruited to telomeric and non-telomeric DNA in response to G4 formation

To address the *in vivo* significance of the CST G4-resolving activity, we next examined whether STN1 subcellular distribution is altered after G4 formation. STN1 localization was examined by indirect immunofluorescence using FLAG antibody and a previously established HeLa cell line expressing FLAG-tagged STN1[21]. G4 was induced by treating the cells with the G4 stabilizing ligands TmPyP4 (50 μM for 24 h) or PDS (10 μM for 24 h). The analysis revealed only a few STN1 foci in DMSO treated control cells whereas TmPyP4 or PDS treatment resulted in a large number of foci. With either drug, the number of cells with 10-20 foci increased ~1.5-fold and the number with >20 foci increased ~2-fold (Fig. 2A-B). To determine whether the STN1 foci were present at telomeres, we performed telomere FISH to monitor the extent of FLAG-STN1 and telomere co-localization. The FISH revealed that ~20% of the STN1 foci were present at telomeres (~17.6% for TmPyP4 and ~23.1% for PDS) (Fig. 2C). These results demonstrate that G4 formation triggers STN1 redistribution and accumulation at specific sites in the nucleus. They further suggested that CST may be recruited to chromatin to aid in the resolution of both telomeric and non-telomeric G4 structures.

**Figure 2.**
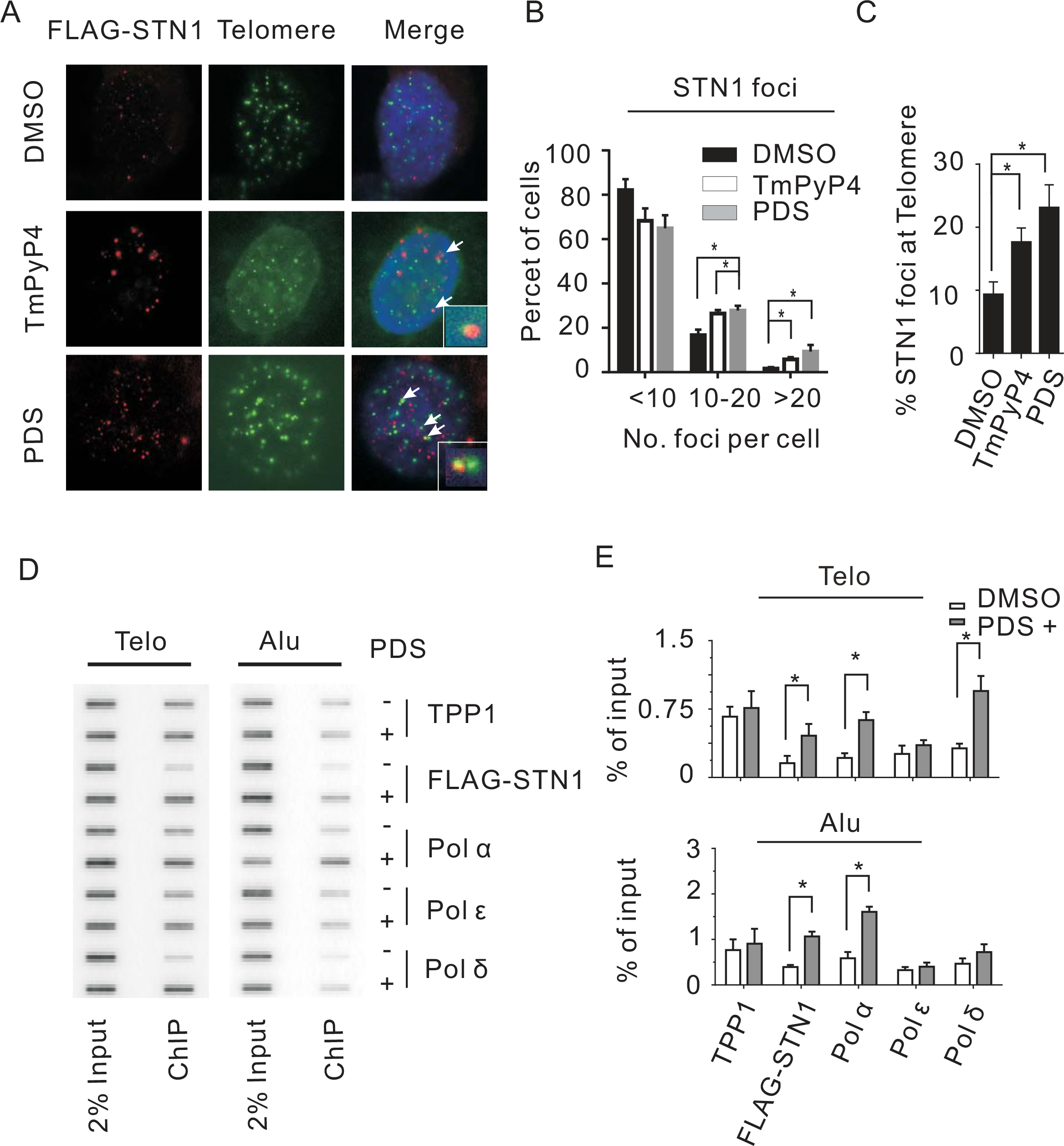
CST is recruited to telomeres after G4 stabilization. (A) Co-localization of FLAG-STN1 and telomeres in HeLa cells after treatment with G4 stabilizers TmPyP4 or PDS. Red, FLAG-STN1 immunostaining; Green, Telomere FISH; Blue, DAPI. (B) Quantification showing FLAG-STN1 foci at telomere induced by G4 ligands. (C) Quantification showing increased numbers of FLAG-STN1 foci following treatment with TmPyP4 or PDS. (C-D) ChIP analysis to examine changes in localization of the indicated proteins to telomeres or Alu repeats after PDS treatment. (D) Slot-blot analysis of DNA precipitated with antibody to TPP1, FLAG (STN1), pol α, pol ε or pol ɛ. Blot was hybridized with a telomere (Telo) or Alu probe. (E) Quantification of ChIP. Telo and Alu ChIP signals were normalized against input DNA. (B & D) Values are mean ± SEM, n = 3 experiments, * P < 0.05. Related to Figure S2.

To directly examine whether G4 stabilization affects CST chromatin association, we used Chromatin Immunoprecipitation (ChIP) assays to quantify changes in CST abundance on telomeric DNA and Alu repeats following 24 hrs PDS or TmPyP4 treatment. The ChIP revealed that both PDS and TmPyP4 cause an increase in the amount of STN1 associated with either sequence and an increase of CTC1 with telomeric DNA (Fig.2D-E & Fig. S2A-B). The increase in telomere localization after drug treatment provides strong support for CST recruitment to chromatin following G4 formation. The increase in CST at Alu repeats is also likely to reflect G4 stabilization as Alu sequences contain runs of Gs and additional G-repeats can be present in the adjacent genomic DNA[40]. Since the human telomere protein TPP1 interacts with CST through CTC1 and STN1[15], we asked whether PDS affects chromatin association of TPP1. However, ChIP revealed no change in TPP1 abundance at telomeres or Alu sequences, indicating that CST-TPP1 interaction is unlikely to be important for G4-induced recruitment of CST.

Given that DNA polymerases are required to re-initiate replication after fork stalling, and that CST interacts directly with DNA Pol α[10, 41], we next asked whether G4 stabilization also affects DNA polymerase association with telomeres or Alu repeats. Cells were again treated with PDS for 24 hrs to allow G4 accumulation and ChIP was used to quantify DNA Pol α, Pol ε and Pol ɛ association. Interestingly, PDS treatment caused a clear increase in DNA Pol α and Pol ε at telomeres but Pol ɛ was largely unchanged. Since Pol α and Pol ε are required for lagging strand replication while Pol ɛ is needed for leading strand replication, this result implies that G4 stabilization affects lagging but not leading strand replication at telomeres. At Alu repeats, PDS treatment only caused a significant increase in Pol α accumulation, raising the possibility of a different outcome with increased primer synthesis by Pol α but no subsequent primer extension by Pol ε or ɛ. Despite this difference, it is notable that G4 stabilizers cause a simultaneous increase in CST and Pol α at telomeres and Alu repeats because this fits with the known role of CST in enhancing priming by Pol α[10, 11]. Overall, our data suggested that CST might prevent G4s from having adverse effects on DNA replication either by resolving the DNA structure and/or stimulating re-priming after fork stalling.

### G4 formation is increased by loss of STN1

Given the ability of CST to bind and unfold G4 DNA *in vitro*[26], we predicted that cellular depletion of CST would result in increased G4 accumulation *in vivo*. To test this prediction, we monitored G4 abundance in previously characterized STN1 knock-down (STN1 sh) and control (non-target, NT sh) cell lines[14, 18]. G4 formation was detected by immunofluorescence using the antibody BG4 raised against human telomeric DNA[42]. Prior to antibody staining, fixed cells were subjected to cytoplasmic extraction and RNase treatment to remove any RNA G4. Additionally, the specificity of the antibody for DNA versus RNA G4 was confirmed by showing that DNase1-treatment of cells removed BG4 staining whereas RNase treatment did not (Fig. S3). As reported previously[42], we observed co-localization of G4 foci with a subset of telomeres in untreated cells. Treatment with TmPyP4 then caused an increase in G4 staining both at telomere and elsewhere in the genome (Fig. 3A-B). When we examined the effect of STN1 knockdown, we also observed a significant increase in the number of G4 foci relative to control cells (Fig. 3A-B), indicating that CST suppresses G4 accumulation. Treatment of the STN1-depleted cells with TmPyP4 caused a further increase in G4 foci both at telomeres and non-telomeric regions. Given that CST can bind G4 DNA, we next asked whether STN1 localizes to G4s *in vivo*. Interestingly, immunostaining revealed that STN1 and G4 foci rarely co-localized (Fig. 3C & Fig. S4 A-C). This finding implies that once CST binds a G4 *in vivo*, it is rapidly unwound or blocks G4 antibody binding. The above results, together with the previously described affinity of CST for G-rich ssDNA, lead us to suggest that CST prevents G4 accumulation in two ways. First CST may bind newly formed ssDNA (e.g. the G-rich lagging strand template during telomere replication) to prevent G4 formation. Second, CST my bind and disrupt previously formed G4 structures (e.g. those formed as a result of transcription[43]). Both aspect of CST function would help prevent blocks to DNA replication.

**Figure 3.**
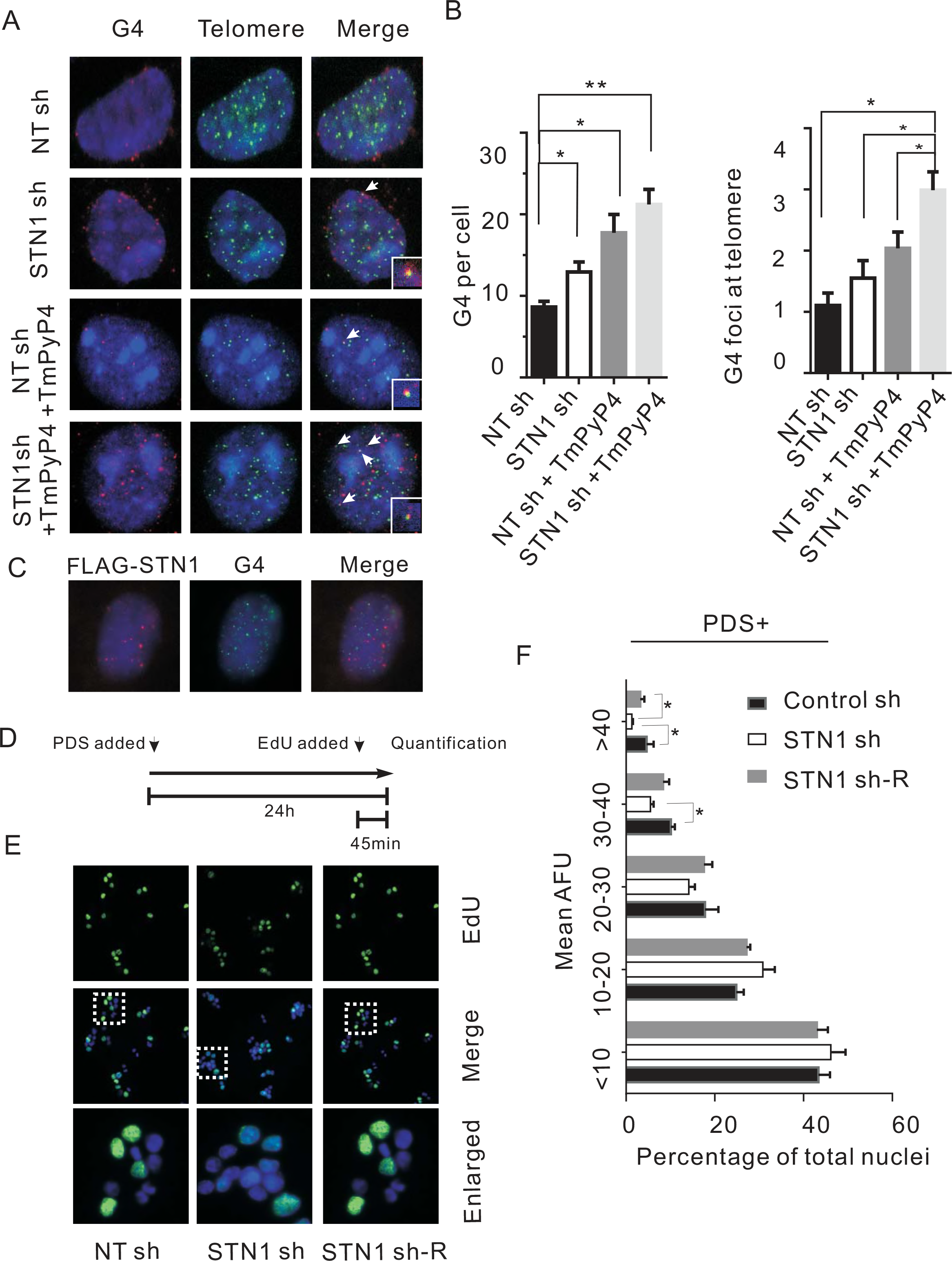
STN1 depletion induces G4 formation and slows DNA replication upon G4 stabilization. (A) Localization of G4 and telomere DNA in control (NT sh) or STN1-depleted (STN sh) HeLa cells. NT, non-target control shRNA. Red, imunolocalization of G4 DNA; Green, telomere FISH; Blue, DAPI. Arrows indicate telomere and G4 co-localization. Arrows indicate telomere and G4 co-localization. (B) Quantification of total (left) and telomeric (right) G4 foci. Values are mean ± SEM, n = 3 experiments, * P < 0.05, ** P < 0.01. (C) Representative image of FLAG-STN1 expressing cells stained with antibody to FLAG and G4 DNA. Red, FLAG-STN1; Green. G4; blue, DNA stained with DAPI. (D) Experimental timeline. NT sh, STN sh or STN1 sh-R cell lines were treated with PDS for 24 hours with EdU added for the last 45 min of treatment. STN1 sh-R, STN1 sh cells expressing sh-resistant FLAG-STN1 allele. (E) EdU incorporation by the indicated PDS-treated cell lines. Blue, DAPI; green, EdU. White boxes indicate regions enlarged in bottom panel. White boxes indicate regions enlarged in bottom panel. (F) Quantification of EdU uptake in the presence of PDS, as measured by mean fluorescence intensity (Mean ± SEM, n = 3 experiments, * P < 0.05). Each bar indicates the percent of nuclei within the indicated AFU range. AFU, arbitrary fluorescence units. Related to Figures S3 to Figures S5.

### STN1 rescues G4-induced inhibition of DNA replication

If CST facilitates DNA replication by removing or preventing G4, loss of CST should result in a slowing of replication and decreased nucleotide uptake. We therefore asked whether STN1 depletion affects EdU uptake during treatment with G4 stabilizers. STN1 sh and control cells were treated with PDS for 24 hrs with EdU added to the culture medium for the last 45 mins of treatment (Fig. 3D). The cells were fixed and the EdU reacted with fluorophore (Fig. 3E). The cells were then imaged and the fluorescence intensity of individual nuclei was quantified. The resulting data are presented both as the mean fluorescence intensity of all nuclei (Fig. S5A) and the percentage individual nuclei that exhibit a specific level of fluorescence (Arbitrary Fluorescence Units, AFU) (Fig. 3F, Fig. S5B). As anticipated, the control cells showed a decrease in the average fluorescence intensity of all nuclei following 24 hrs PDS treatment, indicating that EdU uptake was suppressed by G4 formation (Fig. 3D-F, Fig. S5A-B). The decrease in EdU incorporation likely reflects replication fork stalling at G4 DNA[30, 44]. STN1 depletion caused a further decline in EdU uptake in the PDS-treated cells. This decline was observed both as a decrease in the average fluorescence of all nuclei (Fig. S5A) and a decrease in the percent of nuclei with the highest levels of EdU incorporation (AFU>30) (Fig. 3F). The decline in EdU uptake was largely rescued by overexpression of an sh-resistant FLAG-STN1 allele in the STN1-sh cells[14] (Fig. 3E-F, Fig. S5A-B). Importantly, without PDS treatment, the levels of EdU uptake were similar in the STN1 sh and control cells (Fig. S5A-B) indicating that STN1 depletion in the absence of PDS does not cause a significant difference in the number of cells in S-phase or the rate of replication. We therefore conclude that STN1/CST helps promote passage of the replication fork through genomic regions that are prone to G4 formation. This role in replication most likely stems from the capacity of CST to prevent G4 formation during replication or to unwind previously formed G4. It may also reflect the ability of CST to re-prime replication in situations where fork stalling leads to polymerase dissociation[26].

### Combined STN1 depletion and G4 stabilization induces telomere loss

Since CST functions in multiple aspects of telomere replication, we next examined how G4 stabilization affects telomere integrity in cells that lack STN1. In initial experiments, we examined how the combination of STN1 depletion and PDS treatment affects telomere length. DNA was isolated from control and STN1 sh Hela or U2OS cells after 48 hrs growth with or without PDS, and Telomere Restriction Fragments (TRF) were examined by Southern blot (Fig. 4A and Fig. S6A). The TRF analysis indicated that mean telomere length was not significantly altered by STN1 depletion, consistent with previous reports[6, 7]. Telomere length also remained essentially unchanged by PDS treatment and by PDS treatment combined with STN1 depletion (Fig. 4A-B, Fig. S6A-B). However, the combination of G4 stabilization and STN1 depletion lead to a striking decline in the overall intensity of the telomere hybridization signal. To determine the magnitude of the decline, blots were hybridized with probe to the actin gene to allow normalization for DNA loading. Quantification revealed that the telomere signal decreased by about one third in both HeLa and U2OS cells (Fig. 4C). Given the lack of apparent telomere shortening, this result suggested that a subset of chromosomes had lost essentially all of their telomeric DNA.

**Figure 4.**
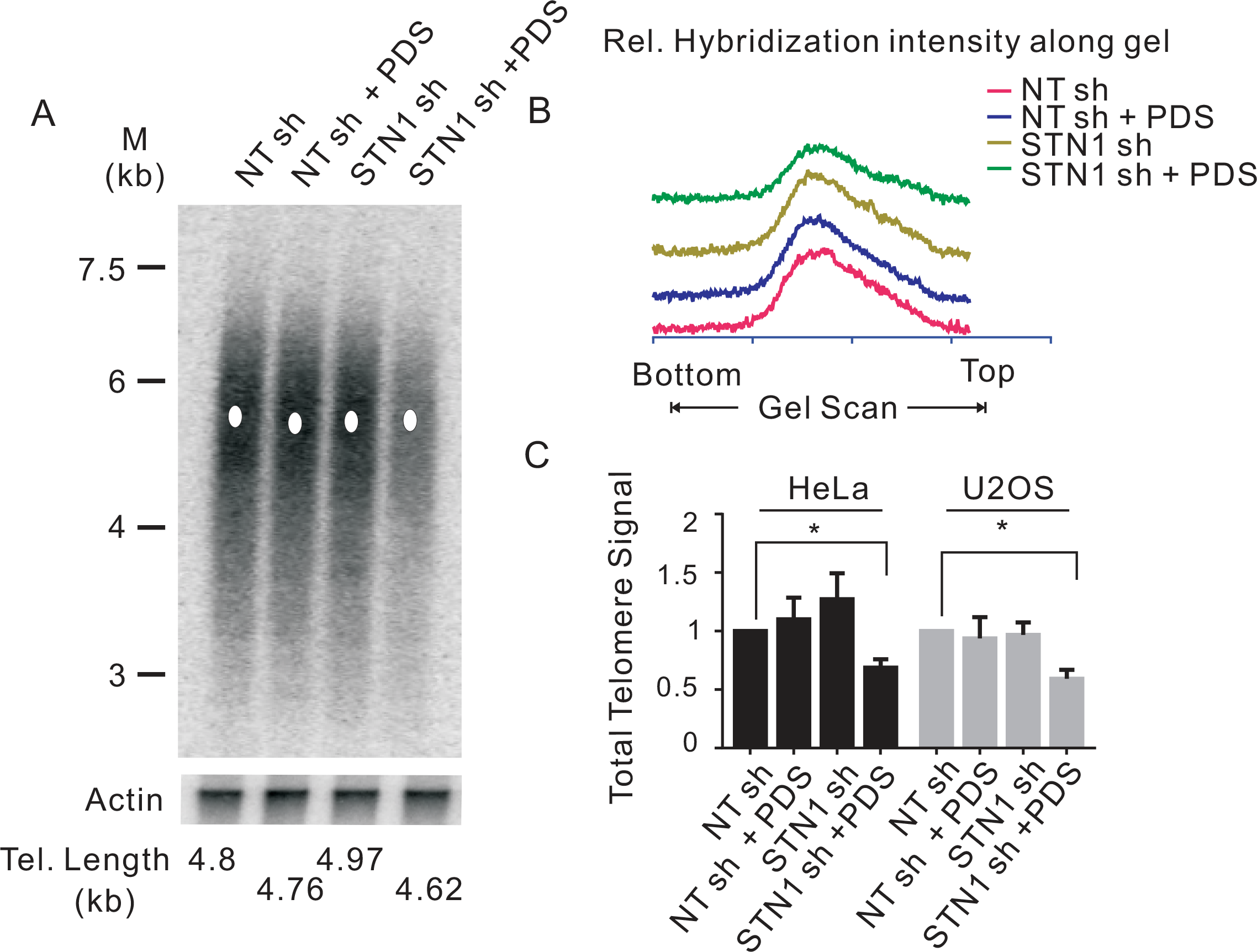
STN1 depletion and G4 stabilization cooperate to cause sudden telomere loss. (A) Southern blot showing telomeric restriction fragments from NT sh and STN1 sh HeLa cells grown with or without 10 μM PDS for 48 hrs. Probe was (T_2_AG_3_)_3_TTA (top) or β-actin (bottom). M, molecular weight markers. (B) Scans showing relative intensity of hybridization signal. Scans for each sample were from top to bottom of blot. (C) Quantification of total telomere hybridization signal from the indicated HeLa and U2OS cell lines grown with or without PDS. Related to Figure S5.

### G4-induced telomere dysfunction is exacerbated by STN1 depletion

CST has previously been shown to facilitate replication through the telomere duplex, with loss of CST leading to slowing of BrdU uptake[18]. CST depletion also leads to two another hallmarks of disturbed telomere replication: the fragile telomere phenotype where individual chromatids exhibit multiple telomere FISH signals (MTS) and sporadic signal free ends (SFE) where telomeres lack detectable FISH signals[14, 17]. Our finding that CST can resolve G4 structure suggested that that CST may facilitate telomere duplex replication by decreasing G4 formation during lagging strand synthesis and/or by preventing G4 accumulation as a result of telomere transcription during TERRA production[45]. Moreover, if CST functions in this manner, the detrimental effects of CST loss on telomere structure should be increased by treatment with G4 stabilizers. To test this prediction, we examined the combined effect of STN1 depletion and TmPyP4 exposure on telomere integrity.

In initial experiments, we prepared metaphase spreads from STN1 sh and NT sh cells grown with and without TmPyP4 for 48 hrs and used telomere FISH to monitor the levels of MTS and SFE. The FISH was performed with probes to both the G-and C-strands so we could determine whether any SFE reflected loss of one or both DNA strands (Fig. 5A). As previously observed[14], quantification of MTS revealed an ~2-fold increase after STN1 knockdown regardless of whether we used the G- or C-strand probe (Fig. 5B-E). When the cells were treated with TmPyP4, the combined STN1 depletion and G4 stabilization lead to a synergistic increase in MTS being detected with either probe. Since G4 stabilization exacerbated the effect of STN1 loss, this result implies that CST averts replication problems by preventing or removing G4 structure. Moreover, the similar frequency of MTS observed with the G- and C-strand probes implies that CST removes G4 structure ahead of the replication fork as fork stalling at a G4 block should affect both daughter telomeres equally.

**Figure 5.**
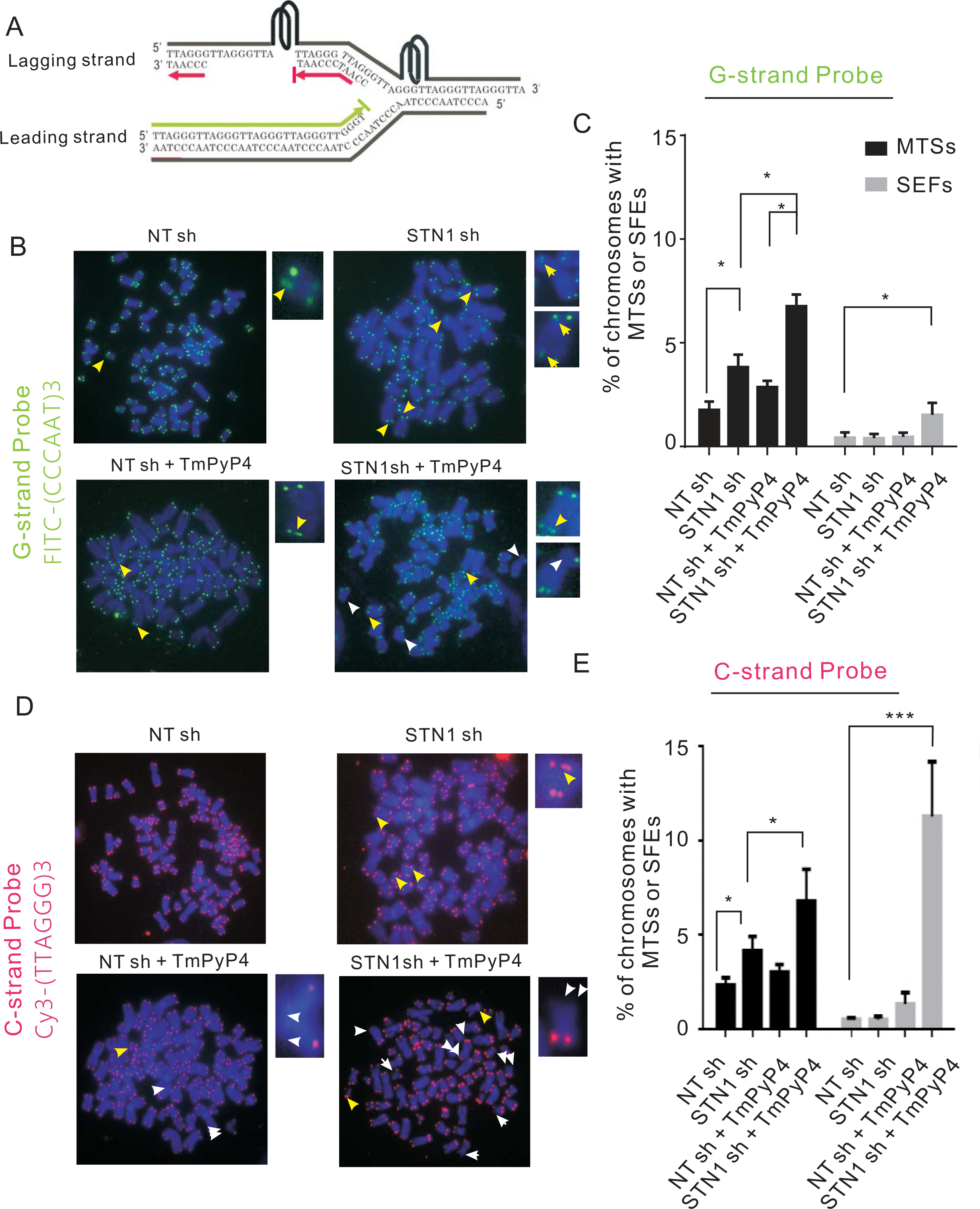
Combined STN1 depletion and G4 stabilization results in preferential loss of telomeric C-strand DNA. (A) Model showing how G4 formation may block DNA synthesis during telomere replication. Red indicates C-strands (identified in telomere FISH using Cy3-TTAGGG)_3_TTA (red) probe), green indicates G-strands (identified using FITC-(CCCAAT)_3_CCC (green) probe). (B-E) Telomere FISH with control NT sh or STN1 sh HeLa cells grown with or without 50 μM TmPyP4 for 48 hr. (B-C) FISH with probe to G-strand. (D-E) FISH with probe to C-strand. (B & D) Representative images of metaphase spreads showing multiple telomere signals (MTS, yellow arrows) and signal free ends (SFE, white arrows). (C & E) Quantification of MTSs and SFEs. (Mean ± SEM, n = 3 experiments, * P < 0.05, * P < 0.01, *** P <0.001).

A remarkably different result was observed when we quantified SFE. As previously reported, STN1 depletion did not increase the frequency of SFE detected using either the G- or the C-strand probe[14]. Treatment of the control NT sh cells with TmPyP4 also had no effect on SFE detected using the G-strand probe but caused a small increase in SFE detected using the C-strand probe. However, when we combined TmPyP4 treatment with STN1 knockdown we saw an ~4-fold increase in SFE with the G-strand probe and an ~16 fold increase with the C-strand probe. Thus, G4 stabilization caused preferential loss of telomeric C-stand DNA. Since leading and lagging strand synthesis are normally tightly coupled, this specific loss of C-strand DNA is unlikely to result from replication fork collapse leading to a DSB as this would not favor loss of one particular strand. However, TmPyP4 treatment is expected to cause G4 accumulation on the parental G-rich strand as a result of ssDNA exposure during Okazaki fragment synthesis. The G4s could then block lagging strand (C-strand) synthesis thus causing loss of C-strand DNA (Fig. 5A) and the increase in SFE detected with the C-strand probe (Fig. 5D-E). In contrast, TmPyP4 should not bind to leading strand replication intermediates and hence the drug is unlikely to cause loss of the parental C-rich strand. We therefore infer that combined STN1 depletion and G4 stabilization causes preferential inhibition of lagging strand replication.

A block in lagging strand synthesis should lead to regions of ssDNA on the parental G-rich strand. This ssDNA would then be expected to trigger DNA damage signals. To test whether combined STN1 depletion and G4 stabilization leads to increased damaging signaling, we used immunostaining with antibody to 53BP1 and the telomere protein TRF2 to monitor the appearance of telomere dysfunction induced DNA damage foci (TIFs). Consistent with previous studies[12], a modest increase in TIFs was observed after STN1 knockdown. Likewise, the TmPyP4 caused a slight increase in TIFs in the control (NT sh) cells. However, combined STN1 depletion and TmPyP4 treatment elicited robust TIFs formation (Fig. 6A-B). This increase in damage signals at chromosome ends that remain bound by TRF2 fits with partial inhibition of lagging strand synthesis as many telomeres would be expected to retain sufficient dsDNA to bind TRF2 while still eliciting a DNA damage response.

**Figure 6.**
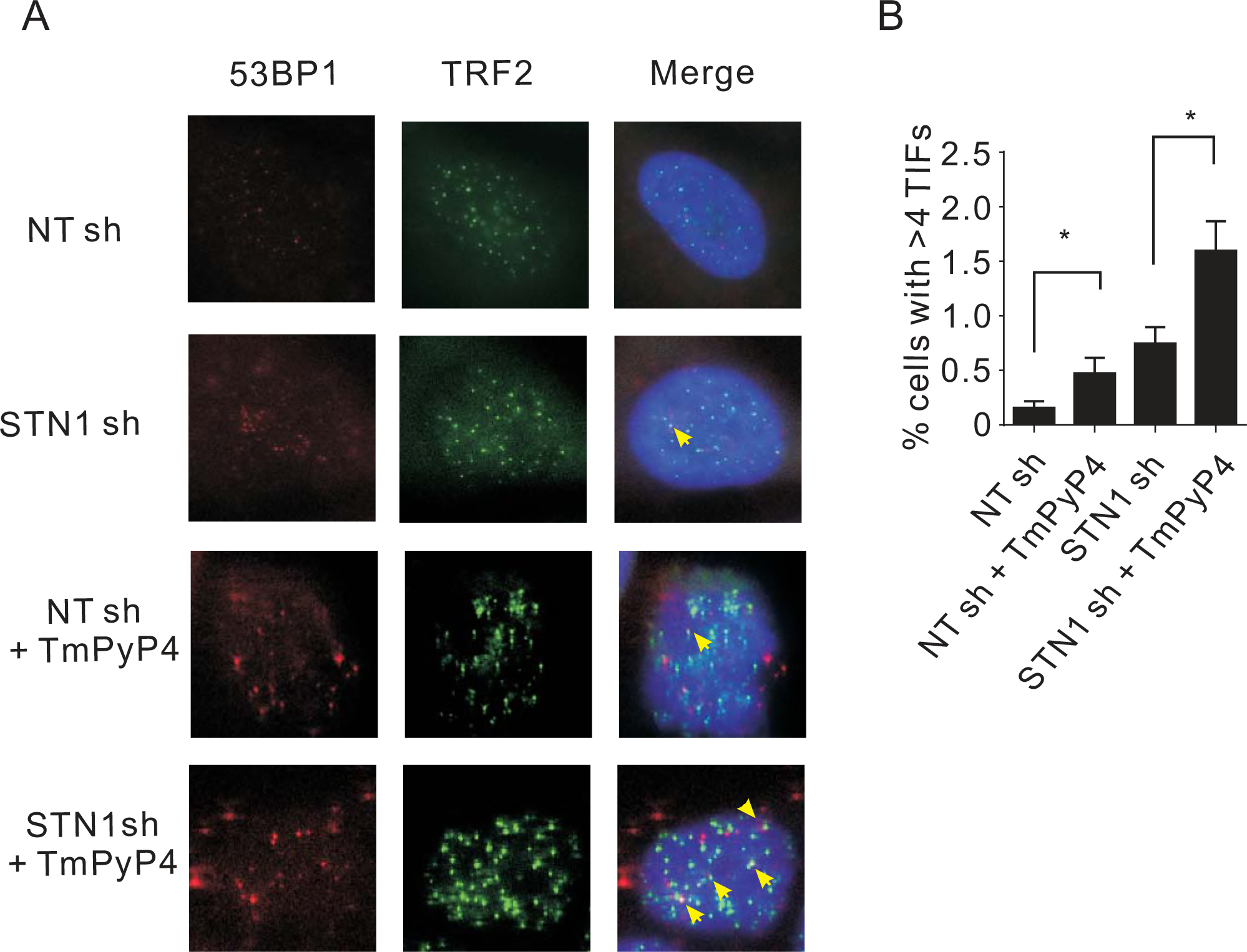
STN1 depletion enhances G4-induced telomere damage. (A) Immunolocalization of 53BP1 (red) and TRF2 (green) in the indicated cell lines grown with or without 50 μM TmPyP4 for 48 hrs. Yellow arrowheads indicate TIFs (sites of 53BP1 colocalization with telomeres). Scale bar, 5 μm. (B) The percentage of cells with ≥4 TIFs was determined for at least 50 cells in each experiment (Mean ± SEM, n = 3 experiments, μ P < 0.05, μμ P < 0.01).

### CST depletion slows lagging strand telomere replication after G4 stabilization

Since our FISH data suggested that combined STN1 depletion and G4 stabilization preferentially inhibited lagging strand telomere replication, we set out to directly test for this by quantifying the fraction of telomeres replicated by leading versus lagging strand synthesis within a set time period. To achieve this, we labeled S-phase cells with BrdU and then used CsCl gradients to separate telomeres newly replicated by leading or lagging strand synthesis (termed leading or lagging telomeres) and quantified the amount of newly replicated telomeric DNA.

HeLa NT sh and STN1 sh clones were synchronized at G1/S with a double-thymidine block, then released into media with or without PDSs and allowed to progress through S-phase for 4 hrs. They were then pulsed labelled with BrdU for 2 hrs and harvested (Fig. 7A & Fig. S7A). DNA was isolated, restriction digested and subjected to CsCl density gradient centrifugation to separate the replicated from unreplicated telomeres (Fig. 7B)[46]. The gradients were fractionated and the relative amount of telomeric DNA in each fraction was determined by slot-blot hybridization using a telomere C-strand probe (Fig. 7C). Telomeres replicated by leading strand synthesis incorporate two BrdU molecules per telomeric repeat (UUAGGG) and hence sediment at a higher density than telomeres replicated by lagging strand synthesis (CCCUAA) and both are separated from any unreplicated telomeric DNA, which has no BrdU incorporation (Fig. 7B).

**Figure 7.**
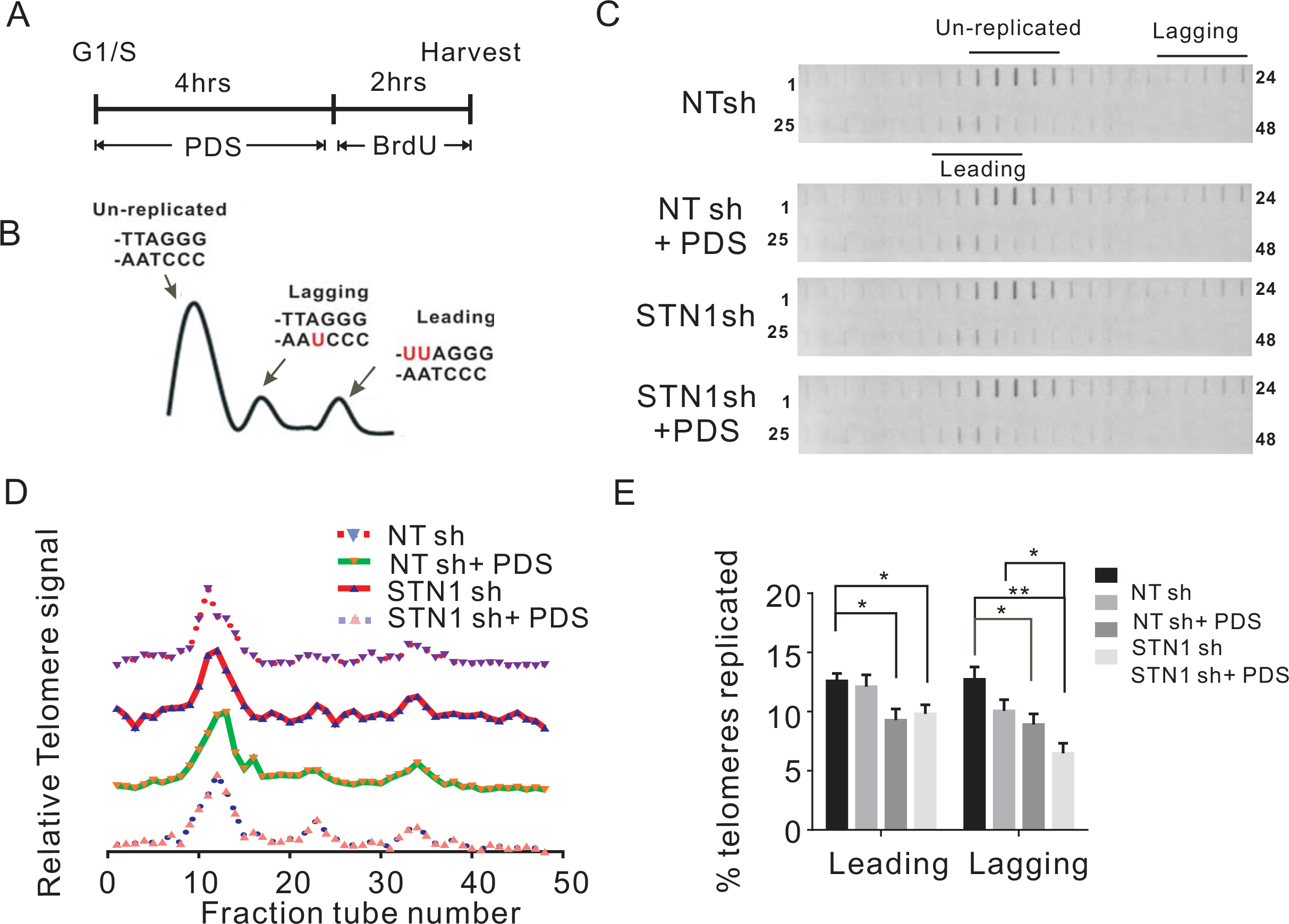
STN1 depletion preferentially decreases lagging strand telomere replication after G4 induction. (A) Experimental timeline. (B) Strategy to separate replicated leading and lagging strand telomeres. Diagram shows BrdU labeling and sedimentation profile for leading, lagging and unreplicated telomeres. Top of the CsCl gradient is to the left. (C) Detection of leading and lagging telomeres by slot blot hybridization with (TTAGGG)_3_TTA probe. (D) Quantification of telomeric DNA hybridization from each gradient fraction. (E) Percent of newly replicated leading and lagging strand telomere signals relative to the total telomere signal. Data are representative of three independent experiments (Mean ± SEM, n = 3, *P < 0.05, **P < 0.01).

Quantification of the hybridization signal from replicated leading and lagging telomeres indicated that PDS treatment decreased replication of lagging but not leading telomeres as might be expected given the G-rich nature of the lagging-strand template (Fig. 7D-E). STN1 depletion decreased the replication of both leading and lagging telomeres suggesting a general slowing of replication through the telomere dsDNA. The combination of STN1 depletion and PDS treatment had no further effect on leading telomere replication. In contrast, lagging telomeres exhibited a significant decrease in newly replicated DNA relative to either single treatment. The preferential effect of the combined treatment on lagging telomeres indicates that CST (STN1) plays an important role in facilitating lagging strand telomere replication in response to G4-induced replication stress.

## Discussion

Although timely resolution of G4 DNA is essential to prevent stalling of DNA replication and ensuing genomic instability, the mechanisms for G4 removal are still not fully understood[30, 47]. Here we identify CST as a novel player in this process. We show that mammalian CST unwinds G4 structure *in vitro*, prevents G4 accumulation *in vivo* and facilitates DNA replication through G4-containing sequence. Although CST is particularly important for telomere replication, its roles in G4 resolution are not limited to telomeres. Induction of G4 structure increases CST localization to both telomeric and non-telomeric chromatin and loss of CST slows bulk genomic DNA replication after G4 stabilization. At telomeres, combined CST depletion and G4 stabilization causes dramatic telomere loss and decreased lagging-strand telomere replication. Thus, the capacity of CST to prevent or remove G4 is essential to maintain telomere integrity. Our findings lead to a model where CST facilitates Okazaki fragment synthesis by resolving or preventing G4 structure on the single-stranded G-rich template DNA. Our data also support a role for CST in preventing replication blocks by removing G4 structures that accumulate ahead of the replication fork.

Failure of lagging-strand synthesis occurs when the exposed template DNA forms secondary structures that block passage of the replicative polymerases[48]. Here we demonstrate the sensitivity of lagging-strand replication to G4 blocks by showing that G4 stabilizers specifically cause the lagging-strand polymerases (Pol α and Pol ε) to accumulate at telomeres. We also show that loss of CST greatly exacerbates problems associated with G4 stabilization to cause G4-induced failure of lagging strand synthesis. The result is a reduction in replicated lagging strand telomeres and loss of telomeric C-strand DNA. These findings provide the first direct evidence that CST facilitates DNA replication by resolving blocks to lagging strand synthesis. Although multiple factors affect telomere replication[49–53], it is most unusual to observe the preferential loss of one strand of telomeric DNA. This is because problems associated with telomere dsDNA replication generally cause replication fork stalling. Subsequent fork collapse leads to a DSB and loss of both DNA strands from one or other sister chromatid[54, 55]. Our finding that CST depletion causes a specific reduction in C-strand DNA implies a lack of replication fork collapse despite the perturbation of lagging strand synthesis. Given that leading and lagging strand synthesis are normally tightly coupled[56], this phenomenon could be explained if initiation of Okazaki fragment synthesis by Pol α remains unaffected. We therefore suggest that G4 formation on the lagging strand template blocks Okazaki fragment extension by DNA Pol α and/or Pol ε rather than the primer initiation step (Fig. 8). The outcome would be incomplete C-strand synthesis (Fig. 5) and accumulation of ssDNA (Fig. 6) without stalling of the replication fork.

**Figure 8.**
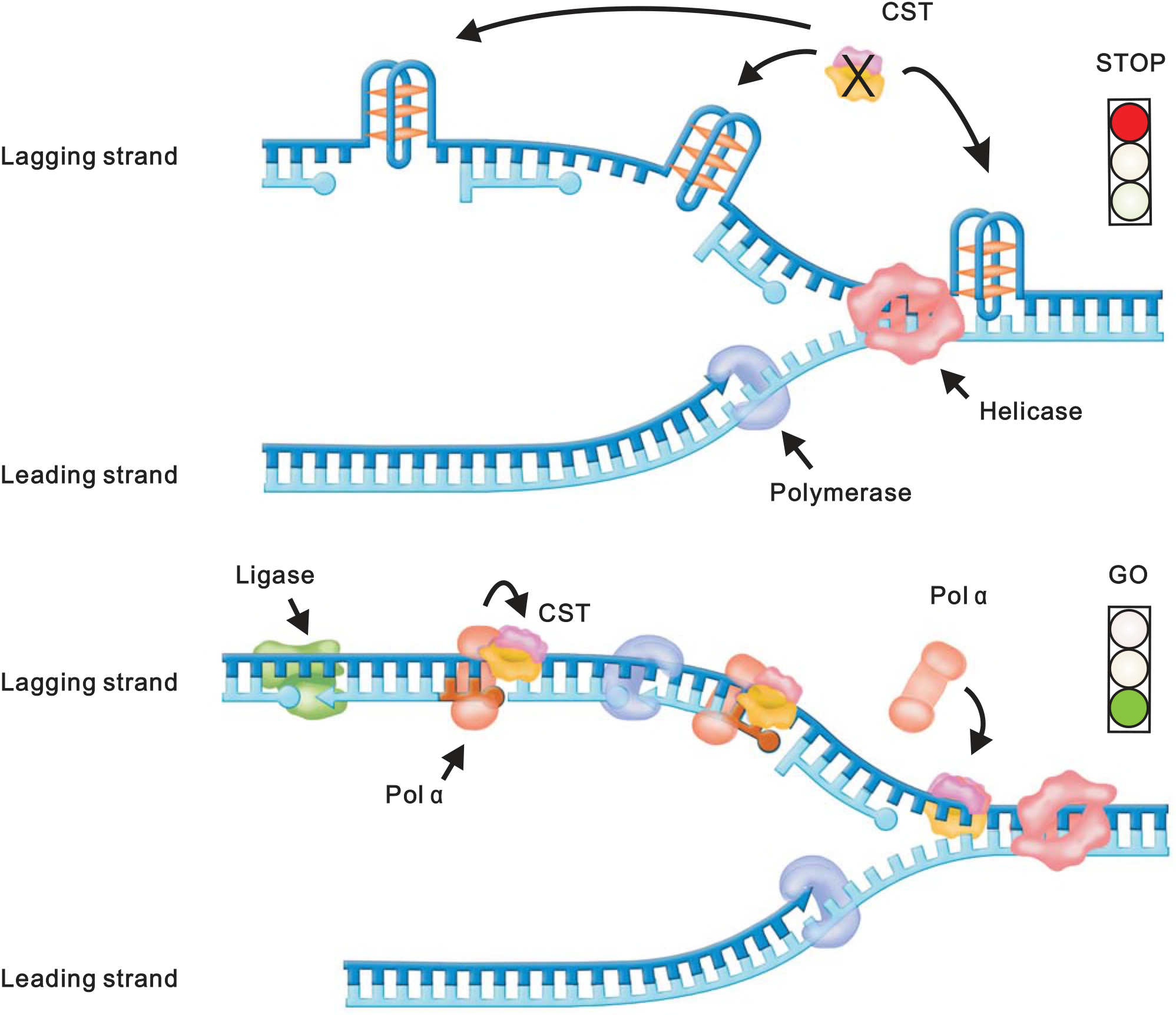
Model for how CST prevents preferential loss of telomeric C-strand DNA following G4 stabilization. Top: In the absence of STN1/CST, replication of the leading strand telomere is unaffected, however formation of G4 structure on the lagging strand template blocks Okazaki fragment synthesis. Bottom: CST/ STN1 resolves or prevents G4 formation. CST may also recruit DNA polα primase to reinitiate replication after G4 resolution.

While the lagging strand template is a prime site for G4 accumulation at telomeres and other G-rich sequences, G4 structures also form during transcription[28, 57]. They tend to occur on the displaced G-rich strands of R-loops and once formed they stabilize the R-loop[58]. The resulting G4/R-loop combination can then block passage of the replication fork. Since, TERRA transcription makes telomeres especially prone to R-loop formation, efficient G4 removal ahead of the replication fork is essential to prevent fork stalling and collapse[59]. Evidence that CST participates in removal of G4 from telomeric dsDNA comes from our finding that combined CST depletion and G4 stabilization causes MTS levels to increase on both leading and lagging telomeres. Since MTS are a hallmark of replication fork stalling[52], this result supports a role for CST in removing G4 structure ahead of the replication fork. Replication fork stalling and collapse into a DSB is also likely to explain the ~1.5% of telomeres that exhibit SFE with loss of G-strand DNA after CST depletion and G4 stabilization (Fig. 5C). Although the current study is focused more heavily on the role of CST in telomere replication, we show that CST depletion also decreases bulk genomic DNA replication after G4 stabilization. We therefore predict that the ability of CST to prevent or resolve G4 will be important for efficient DNA replication at G-rich regions throughout the genome.

The detrimental effects of G4 on DNA replication and genome stability, likely explains why cells have evolved a wide range of factors to remove these structures (e.g. ssDNA binding proteins, helicases and nucleases)[60–63]. Depending on the context of the G4, specific factors may be better suited for G4 prevention or resolution. For example, RPA and CST have similar ssDNA binding and G4 unfolding activities[64]. However, use of CST to resolve G4s during DNA replication may be advantageous because CST can unfold G4 structures more rapidly than RPA and CST-coated ssDNA does not cause ATR activation (Fig. 1 and [19]). Helicases are another major group of proteins known to promote efficient resolution of G4 structure[65]. However, engagement of a helicase on G4-containing DNA requires either an adjacent region of ssDNA for loading or interaction with another protein. For example, TRF1 is needed to recruit the BLM helicase to telomeres for G4 resolution[66]. In contrast to helicases, CST can load bind to and unwind G4 in the absence of adjacent ssDNA[26].

Going forward it will be important to explore how CST interfaces with other G4 resolving activities in a cell. It could be that CST serves as a first line of defense against G4-induced replication issues by coating the exposed ssDNA to prevent G4 formation without activating DNA damage signaling. CST-mediated G4 regulation may be also important for processes beyond DNA replication. G4 structures are expected to form at thousands of sites in the genome where they can affect a wide ranges of cellular processes including transcription and translation. Interestingly, mutations in CST cause Coats plus, a lethal disease with very pleiotrophic symptoms[67–69]. While some symptoms mirror those of telomere maintenance syndromes such as dyskeratosis congenita, others are unique. It is possible that the diverse symptoms of Coats plus patients reflect both altered gene expression and genome-wide replication issues due to loss of normal CST G4-resolving activity.

## Material and Methods

### Cell culture

Different HeLa, HeLa 1.2.11 cell clones and U2OS cells were cultured in RPMI 1640, HCT116 cells in McCoy’s 5A media containing 10% FBS, antibiotics and glutamine.

### EMSA assays

For Kd analysis, CST (0.75-60 nM) was incubated with 32P-labeled Tel21 (0.05 nM) in binding buffer (10 mM Tris pH 7.4, 1 mM DTT) containing either150 mM NaCl or 150mM LiCl for 2h at 4°C to reach reaction equilibrium. DNA binding was then monitored by EMSA. Samples were separated in 0.7% agarose gels with 1 × TAE for 1 h at 90V and 4°C and then quantified by PhosphorImaging. To determine Kd, app, the amount of bound versus free DNA was quantified using ImageQuantTL software. Data were fit to a one site specific saturation binding equation using Graphpad prism software.

### CD Spectra

Tel21 DNA (0.5 μM) were dissolved in 10 mM Tris (pH 7.4) with 150 mM NaCl or 150 mM LiCl, heated at 95 °C for 5 min, and then cooled down to room temperature over night.CD spectrum was collected from 320 to 220 nm at 1-nm bandwidth on a Chirascan Plus CD spectropolarimeter (Applied Photophysics) with a 10-mm pathlength at 25°C. Buffer blank correction was made for both samples.

### Fluorescence Resonance Energy Transfer

Tel21 DNA labeled with fluorescein (FAM) at the 5’-end and tetramethylrhodamine (TAMRA) at the 3’-end, were purchased from Takara Biotech (Dalian, China).) The DNA (1 μM) was incubated with 25 or 100 nM proteins in 10 mM Tris (pH 7.4) buffer containing 150 mM NaCl for 2h at 4°C. Then the fluorescence measurements were carried out on a Spex Fluorolog-3 spectrofluorometer (HORIBA Jobin Yvon, France) at 25°C. The excitation and emission slits were both 5 nm. Excitation was set at 480 nm, and emission was collected from 490 to 650 nm. Lamp fluctuations were corrected by using the reference channel. A buffer blank was subtracted for all spectra.

### Immunofluorescence (IF)

Cells were grown in chamber slides and fixed in 4% paraformaldehyde for 15 minutes, washed 3 times with PBS, permeabilized with 0.15% Triton X-100 for 15 minutes, washed 3 times for 5 minutes with PBS, blocked with 10% BSA at 37o C for 1 hr in humidified chamber, incubated with primary antibodies for overnight at 4o C, washed with PBS three times, incubated with secondary antibody at r.t. for 1 hr, and finally washed 3 times in PBS. Slides were treated with cold ethanol series, dried in the dark, and DAPI-containing mounting medium (Vector Labs) was applied for visualization. Images were taken under Nikon ECLIPSE Ti fluorescence microscope with a 100X objective. Antibodies used were as follows: Flag-tag, Sigma-Aldrich, F3165; TRF2, Millipore, 05-521; 53BP1, NOVUSBIO, NB100-304; HA-tag, Immunoway, YM3003; Dylight 488-anti-mouse IgG (ThermoFisher) and Dylight 549-anti-rabbit IgG (ThermoFisher).

### Chromatin Immunoprecipitation

As previously described[19], cells were fixed with 1% formaldehyde for 20 min. then treated with 200 mM glycine for 10 min to quench the reaction. Cell were pelleted by centrifugation, suspended in swelling buffer (25 mM HEPES, pH 7.9, 10 mM KCl, 1.5 mM MgCl2, 1 mM EDTA, 1 mM DTT, 0.25% Triton X-100 and protease inhibitors) on ice for 10 min, pelleted and incubated in sonication buffer (50 mM HEPES, pH 7.9, 150 mM NaCl, 1 mM EDTA, 0.1% Sodium deoxycholate, 0.1% SDS, 1% Triton X-100 and protease inhibitors) and sonicated for 20 min. in a sonication system (Dajin). Samples were centrifuged at 14 000 rpm for 10 min and the supernatant used for ChIP. Samples containing supernatant (0.3 mg protein), antibody (10 μg TPP1, abcam, ab54685; 5 μg Flag, Sigma-Aldrich, F3165; 15 μg Pol α, Santa Cruz, sc-5921; 15 μg Pol ε, Santa Cruz, sc-10784; 15 μg Pol ɛ, Santa Cruz, sc-398582) and 1 μg yeast RNA were incubated overnight at 4°C. Protein A/G PLUS agarose beads (Santa Cruz) were then added and samples incubated for 1 h at 4°C. Beads were washed sequentially with wash buffer A (20 mM Tris-HCl, pH 8.0, 1 mM EDTA, 0.1% SDS, 1% Triton X-100, 150 mM NaCl), buffer B (20 mM Tris-HCl, pH 8.0, 1 mM EDTA, 0.1% SDS, 1% Triton X-100, 500 mM NaCl), buffer C (20 mM Tris-HCl, pH 8.0, 1 mM EDTA, 0.5% NP40, 0.5% sodium deoxycholate, 250 mM LiCl) and TE buffer (10 mM Tris-HCl, pH 8.0, 1 mM EDTA). The immunoprecipitate was eluted in 450 μl elution buffer (1% SDS, 0.1 M NaHCO3), and cross-linking was reversed by incubation at 65°C overnight. The eluate was brought to 10 mM EDTA, 40 mM Tris-HCl, pH 6.8 and treated with RNase A at 37°C for 1 h and protease K at 55°C for another hour and the DNA purified by phenol-chloroform extraction. The input and precipitated DNAs were analyzed by slot blot hybridization with either Tel or Alu probes and the signal quantified by Phosphorimager. The background from the no antibody control was subtracted and the amount of precipitated DNA was calculated as a percentage of the corresponding input.

### Telomere FISH

FISH was performed on metaphase spreads from methanol/acetic acid fixed cells as previously described (18), with FITC G-strand probe (5’ CCCTAACCCTAACCCTAA, Biosynthesis) or TelG-Cy3 PNA C-strand probe (5’ GGGTTAGGGTTAGGGTTA, Biosynthesis). Images were taken at a constant exposure time. Multiple Telomere Signals (MTS) and Signal free ends (SFE) are quantified by eye.

### Telomere Southern Blot

Purified genomic DNAs were restriction digested then run on 1% agarose gels. Then gels were denatured, dried and hybridized with ^32^P-labeled (TAAGGG)_3_TTA probe to the telomere. The signal was quantified by Phosphorimager. Signal was quantified by PhosphorImager, and mean telomere length was determined by dividing each lane into 100 boxes using ImageQuant and applying the formula ∑Sig/∑(SigI/LI), where Sig is the sum of the signal from all 100 boxes, SigI is the signal in an individual box, and LI corresponds to the average length of the DNA in that box as determined using DNA markers and a standard curve. To determine the relative amount of telomeric DNA in each sample, the gel was then hybridized with a probe to the β-actin gene and the signal quantified. The total telomere signal from individual lanes was divided by the signal from the β-actin gene to normalize for gel loading.

### Analysis of Genomic DNA and Telomere Replication Rates

Whole genome replication rates were determined based on EdU uptake[14]. Cells were cultured with or without PDS for 24 hours, then pulsed with 50 μM EdU for 45 min. Cells were stained for EdU uptake using Click-It AlexaFluor488 according to the manufacturer’s instructions. To quantify EdU uptake, the average fluorescence (AFU) of individual nuclei was determined with Image J using particle analysis with watershed. The defined regions of interest (ROI) were overlaid onto the image with the EdU signal. The mean AFU was then acquired for each ROI. These numbers were used to determine the average AFU of all nuclei and to show the distribution of nuclei with different ranges of AFU/EdU uptake. At least 700 or 400 nuclei were scored for each independent experiment.

For telomere replication analysis, HeLa cells were released into S phase after a double-thymidine block and cultured in the media containing 50 μM PDS for 4 hrs, then cells were washed and pulse labeled with BrdU (100 mM) for 2 hrs. Genomic DNA was isolated by high-salt precipitation. Leading and lagging strand daughter telomeres were separated in CsCl density gradients as described[18]. Gradients were fractionated and the amount of telomeric DNA in each fraction was determined by slot blot using ^32^-P labeled C-strand probe. The percentage of newly synthesized telomere was calculated by dividing the replicated leading or lagging strand peak by the sum of all peaks.

## Acknowledgements

We thank Dr. Shankar Balasubramanian (Univerisity of Cambridge, UK) for providing BG4 antibody (plasmid). This work was supported by Ministry of Science and Technology (2017YFC1001904), National Natural Science Foundation of China (91649102, 31771520, 21177091, 21647008, 31471293, 81501386, 81671054, 81771135), the Key Project of Tianjin Research Program of Application Foundation and Advanced Technology (15JCZDJC35100). CMP was supported by National Institutes of Health grant RO1041803.

## Author contributions

YL designed and performed the ChIP assay, G4 staining and contributed to other experiments. YNX and TFL designed and executed the MTS and TIF analysis. They also contributed to other experiments. MMZ performed the EdU incorporation analysis and contributed to other experiments. BW performed the CsCl gradient experiment. QL performed the southern blot. FW designed and directed the project and prepared the manuscript. CMP contributed to project design and manuscript preparation.

---

**Figure S1.**
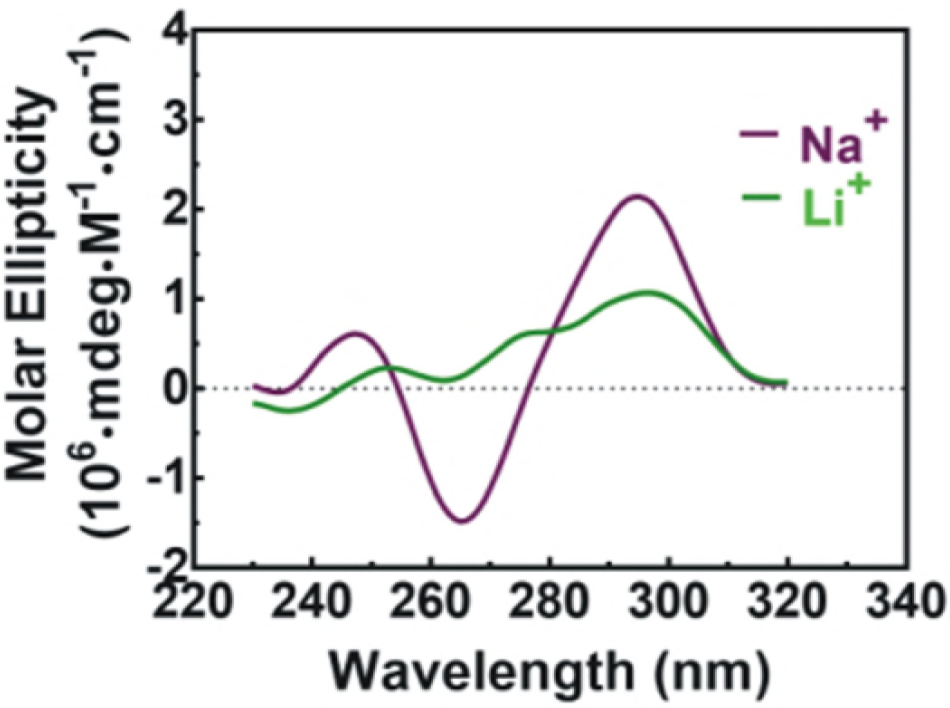
CD spectra of Tel21. The DNA was incubated in 150 mM NaCl (Purple line) or 150 mM LiCl (Green line). The data was obtained with a 0.5 μM strand concentration (pH 7.4) at 25°C. Buffer blank correction was made for both samples.

**Figure S2.**
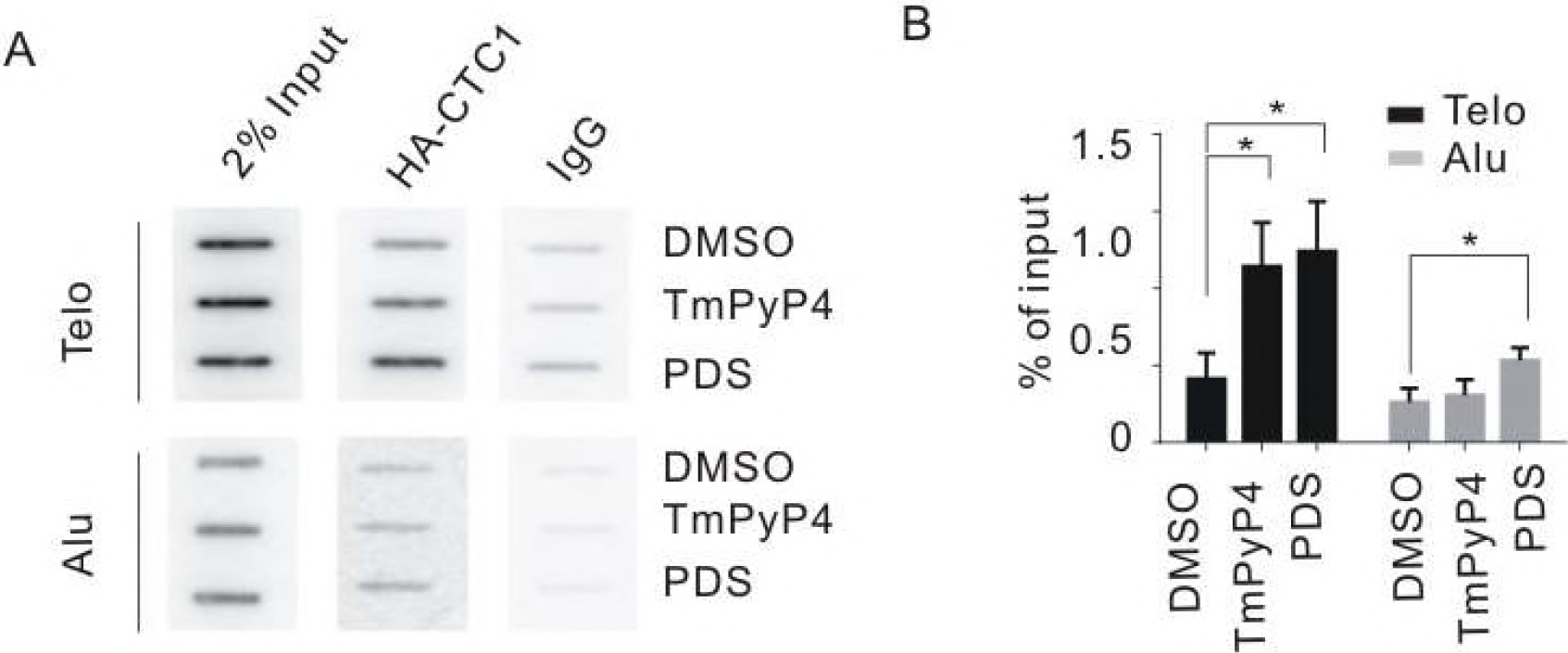
CTC1 is recruited to telomeres upon G4 stabilization. ChIP was performed with HA antibody using HA-CTC1 expressing HeLa cells with and without treatment with TmPyP4 or PDS. (A) Slot blot analysis of precipitated DNA. Hybridization was with telomere probe (Telo) or Alu probe. (B) Quantification of ChIP data. Telo and Alu ChIP signals were normalized against input DNA. Data are expressed as mean ± SEM, n = 3, *P < 0.05. Related to Figure 2.

**Figure S3.**
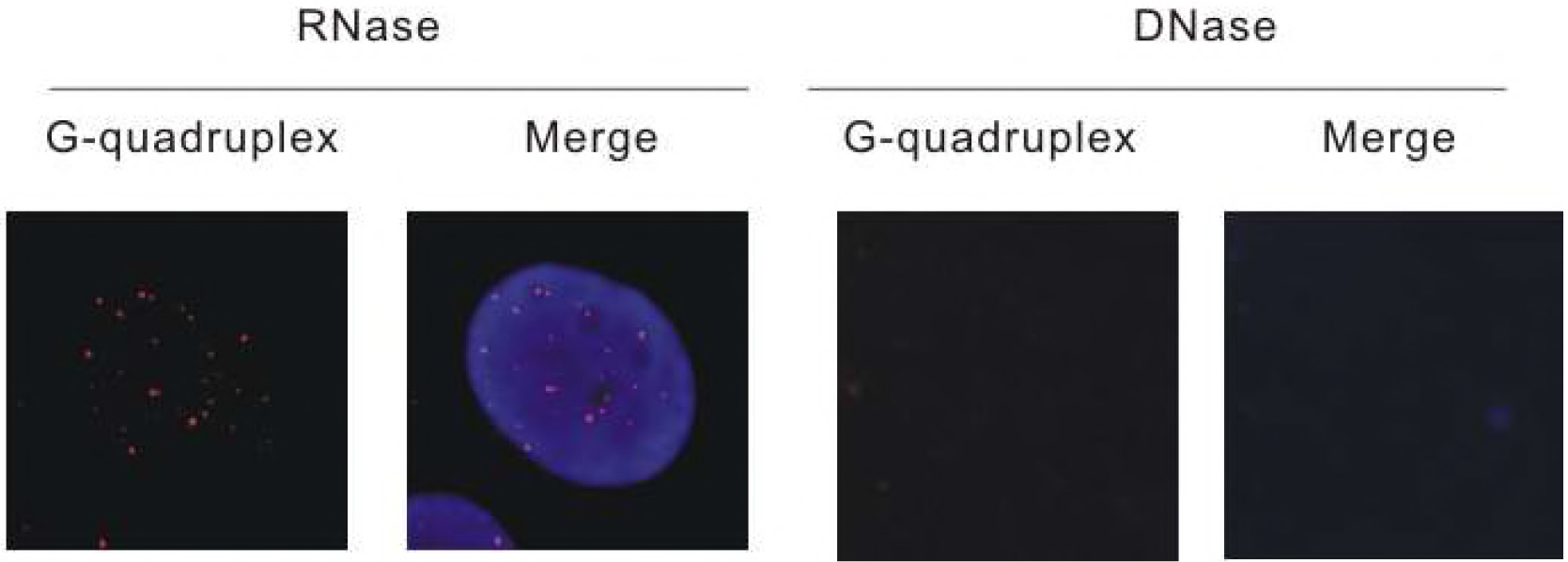
BG4 antibody recognizes DNA but not RNA-based G4 DNA. (A) Immunofluorescence detection of G4 (red) in HeLa cells using BG4 antibody. Cells were treated with RNase (left) or DNase (right) prior to immunolocalization and counterstained with DAPI (blue).

**Figure S4.**
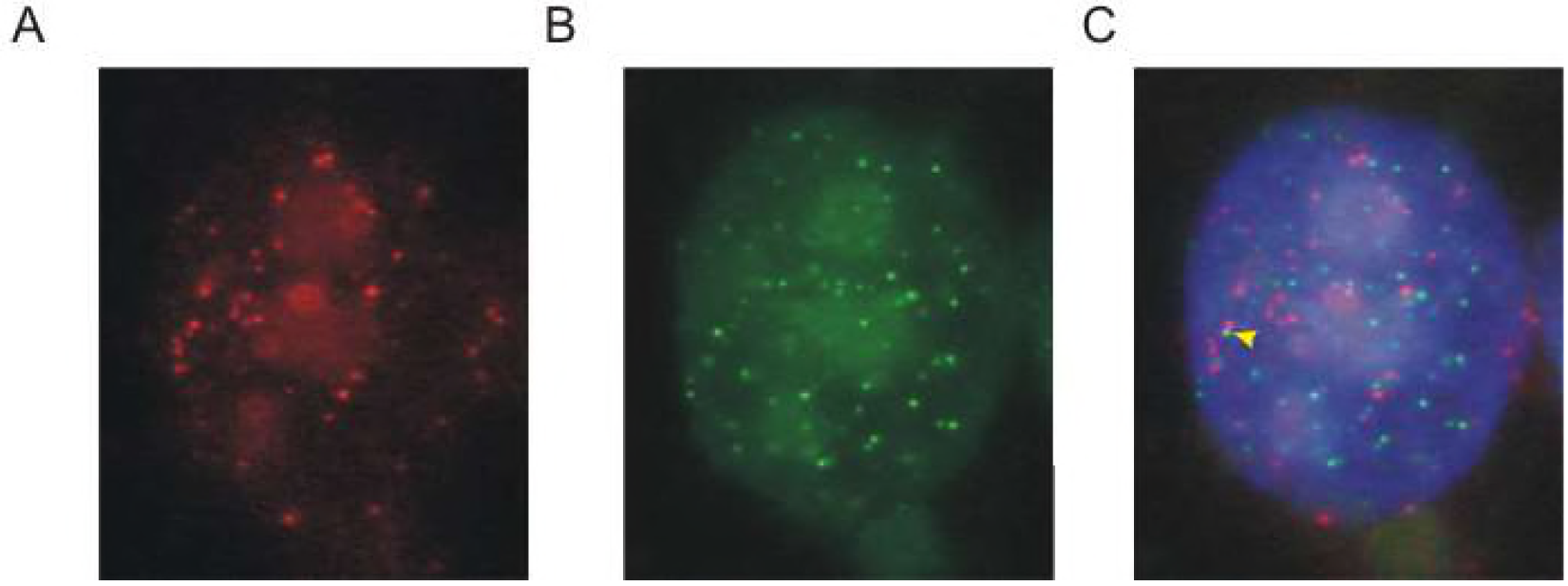
Representative enlarged image of FLAG-STN1 expressing cells stained with antibody to FLAG and G4. Red, FLAG-STN1; Green. G4; blue, DNA stained with DAPI. Related to Figure 3 Yellow arrowhead indicates rare co-localization of FLAG-STN1 and G4.

**Figure S5.**
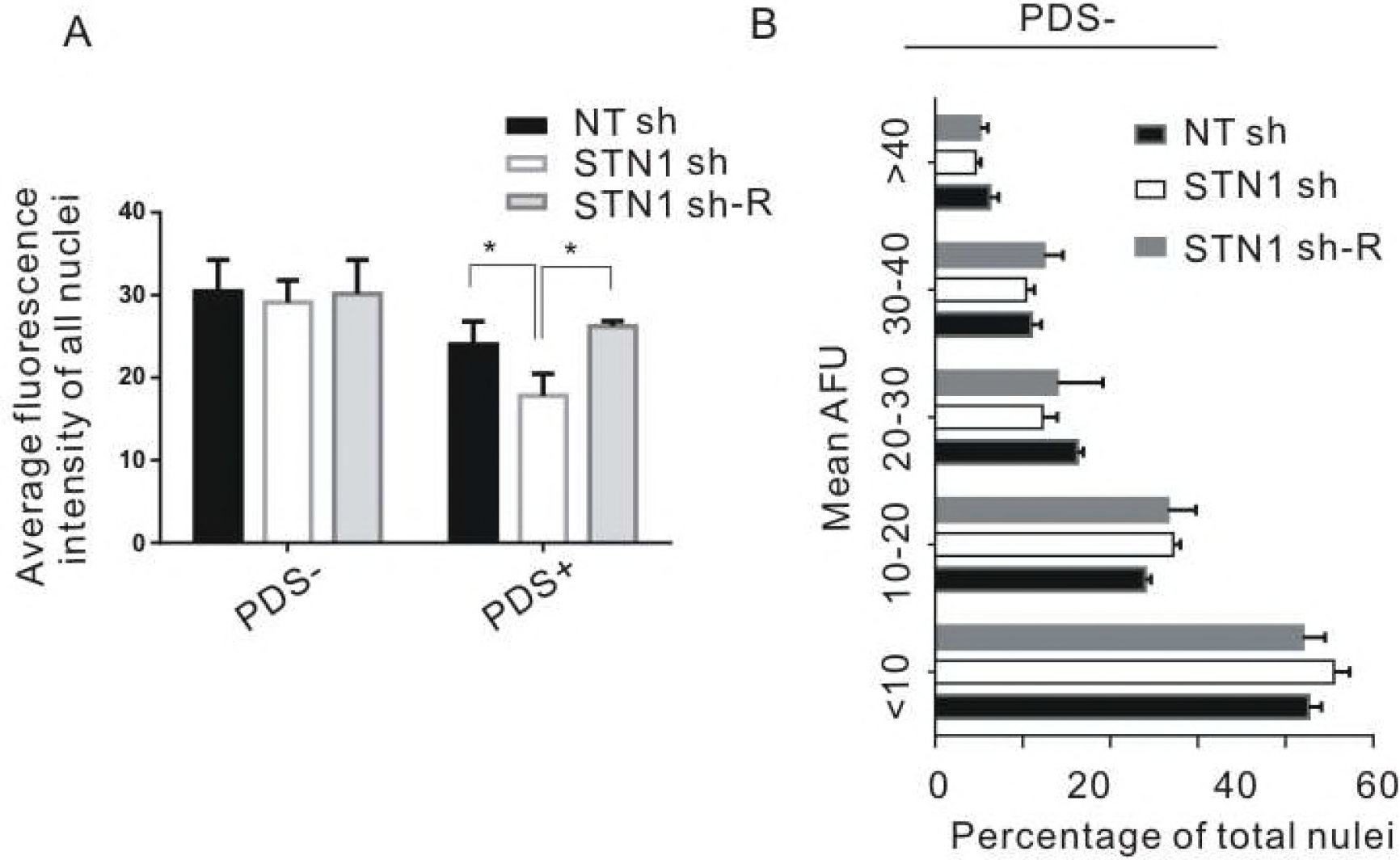
STN1 depletion slows bulk genomic DNA replication after G4 stabilization. (A & B) Quantification of EdU uptake based on fluorescence intensity. (A) Average fluorescence intensity values for all nuclei. The indicated HeLa cell lines were grown with/without 10 μM PDS for 24 hrs. NT sh, non target control; STN1 sh, cells depleted of STN1 with shRNA; STN1 sh-R, STN1 sh cells expressing sh-resistant FLAG-STN1. (B) Quantification of the percent of nuclei within the indicated fluorescence range. EdU labeling was in the absence of PDS treatment. AFU, arbitrary fluorescence units. Data are expressed as mean ± SEM, n = 3, *P < 0.05. Related to Figure 3.

**Figure S6.**
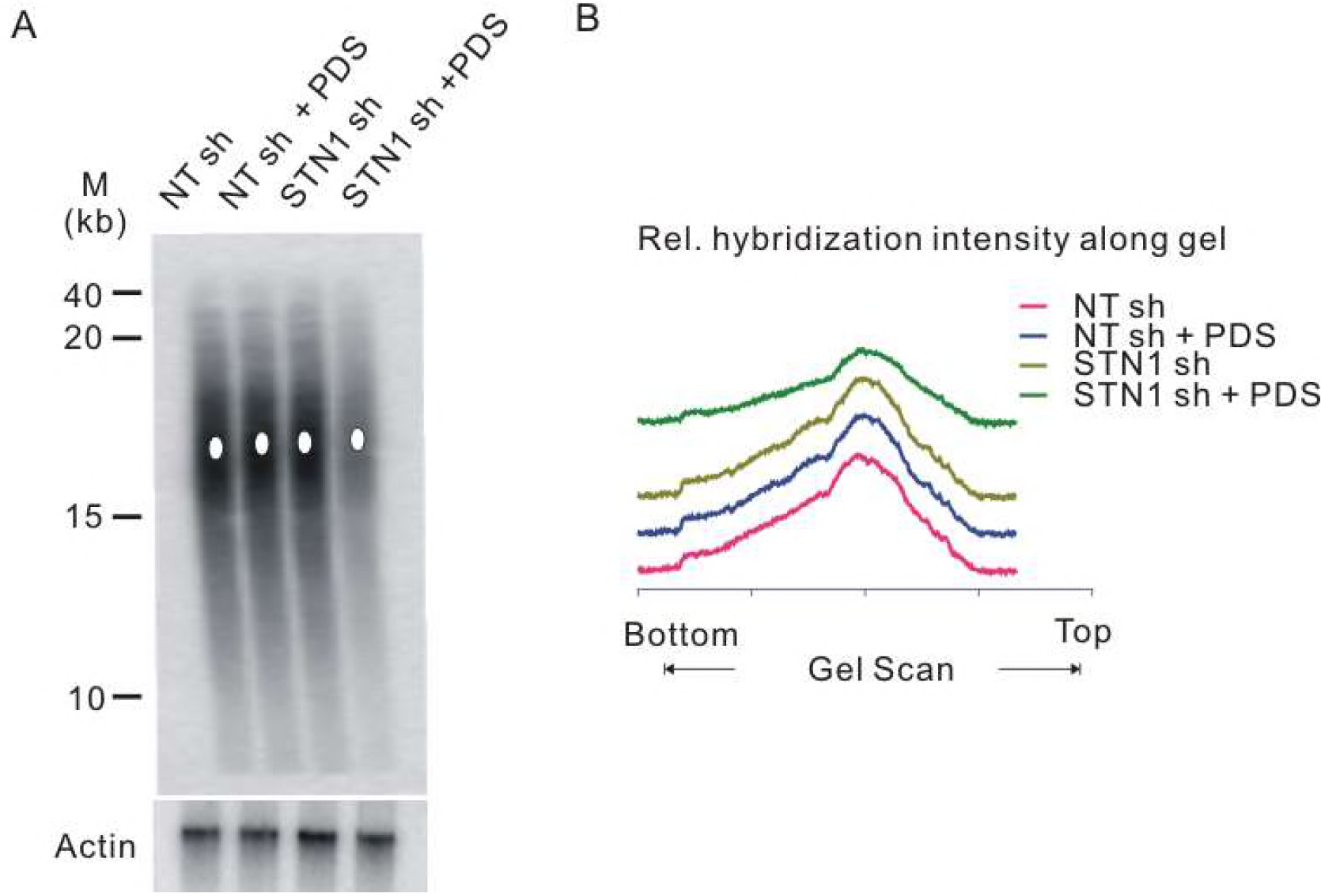
STN1 depletion induces telomere loss not telomere shortening after G4 stabilization. (A) Genomic DNA from pools of U2OS cells (control with NT shRNA or knockdown with STN1 shRNA) was resolved by agarose gel electrophoresis. Telomere restriction fragments were detected with ^32^P-labeled (TTAGGG)_3_TTA telomere probe. M, molecular weight markers. (B) Scans showing relative intensity of hybridization signal. Scans for each sample were from top to bottom of blot. Related to Figure 4.

**Figure S7.**
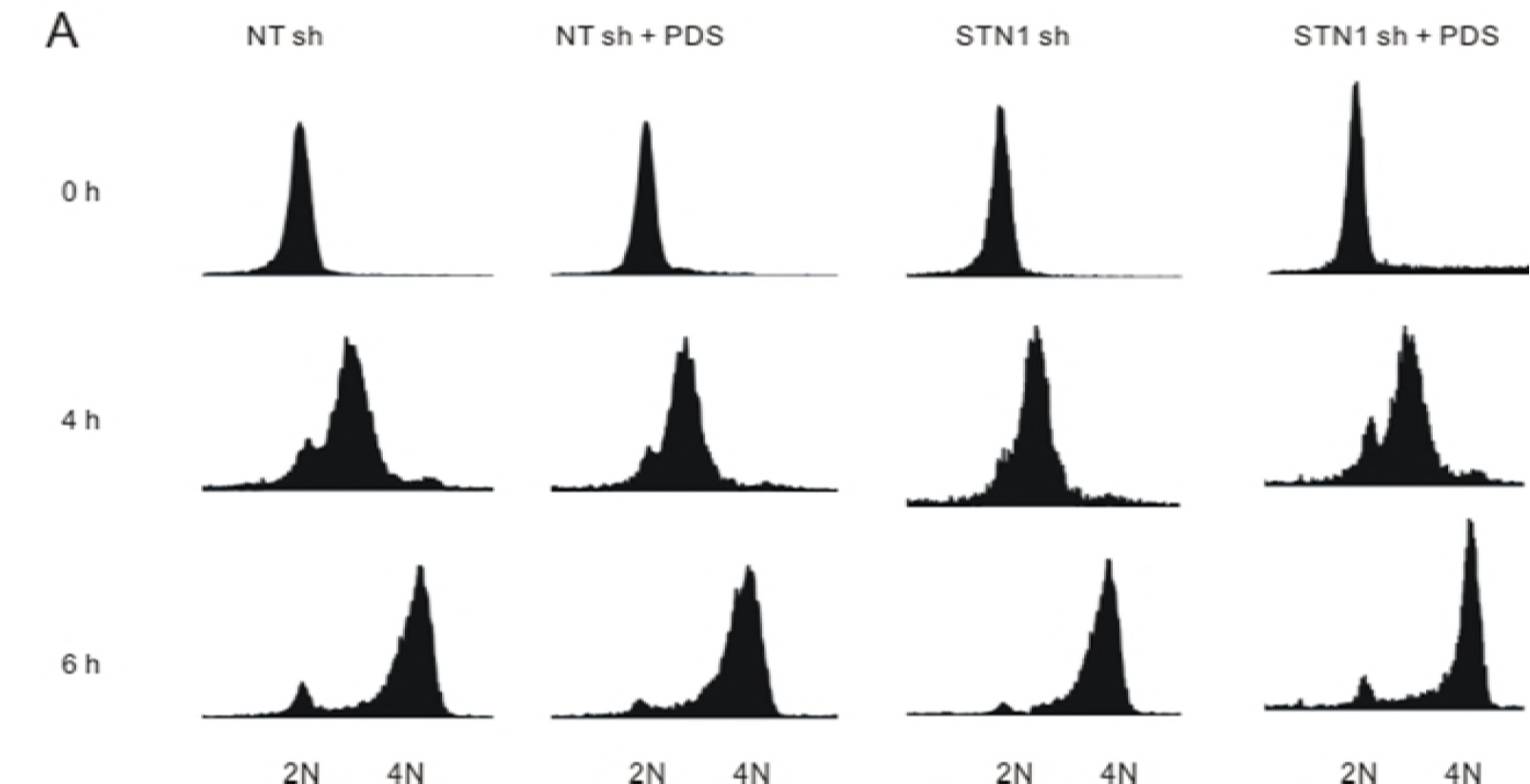
Synchronization of cells used to separate leading and lagging strand telomeres. (A) FACS analysis showing synchrony of cells used to collect DNA in Figure 7D.

## References

1. O’Sullivan, R.J. and J. Karlseder, Telomeres: protecting chromosomes against genome instability. Nat Rev Mol Cell Biol, 2010. 11(3): p. 171–81.

2. Arnoult, N. and J. Karlseder, Complex interactions between the DNA-damage response and mammalian telomeres. Nat Struct Mol Biol, 2015. 22(11): p. 859–66.

3. Stewart, J.A., et al., Maintaining the end: roles of telomere proteins in end-protection, telomere replication and length regulation. Mutat Res, 2012. 730(1-2): p. 12–9.

4. Sfeir, A. and T. de Lange, Removal of shelterin reveals the telomere end-protection problem. Science, 2012. 336(6081): p. 593–7.

5. Kim, J.K., et al., Structural Basis for Shelterin Bridge Assembly. Mol Cell, 2017. 68(4): p. 698–714 e5.

6. Miyake, Y., et al., RPA-like mammalian Ctc1-Stn1-Ten1 complex binds to single-stranded DNA and protects telomeres independently of the Pot1 pathway. Mol Cell, 2009. 36(2): p. 193–206.

7. Surovtseva, Y.V., et al., Conserved telomere maintenance component 1 interacts with STN1 and maintains chromosome ends in higher eukaryotes. Mol Cell, 2009. 36(2): p. 207–18.

8. Wellinger, R.J., The CST complex and telomere maintenance: the exception becomes the rule. Mol Cell, 2009. 36(2): p. 168–9.

9. Ruggiero, E. and S.N. Richter, G-quadruplexes and G-quadruplex ligands: targets and tools in antiviral therapy. Nucleic Acids Res, 2018.

10. Goulian, M. and C.J. Heard, The mechanism of action of an accessory protein for DNA polymerase alpha/primase. J Biol Chem, 1990. 265(22): p. 13231–9.

11. Ganduri, S. and N.F. Lue, STN1-POLA2 interaction provides a basis for primase-pol alpha stimulation by human STN1. Nucleic Acids Res, 2017. 45(16): p. 9455–9466.

12. Huang, C., X. Dai, and W. Chai, Human Stn1 protects telomere integrity by promoting efficient lagging-strand synthesis at telomeres and mediating C-strand fill-in. Cell Res, 2012. 22(12): p. 1681–95.

13. Gu, P., et al., CTC1 deletion results in defective telomere replication, leading to catastrophic telomere loss and stem cell exhaustion. EMBO J, 2012. 31(10): p. 2309–21.

14. Stewart, J.A., et al., Human CST promotes telomere duplex replication and general replication restart after fork stalling. EMBO J, 2012. 31(17): p. 3537–49.

15. Chen, L.Y., S. Redon, and J. Lingner, The human CST complex is a terminator of telomerase activity. Nature, 2012. 488(7412): p. 540–4.

16. Feng, X., et al., CTC1-STN1 terminates telomerase while STN1-TEN1 enables C-strand synthesis during telomere replication in colon cancer cells. Nat Commun, In Press.

17. Kasbek, C., F. Wang, and C.M. Price, Human TEN1 maintains telomere integrity and functions in genome-wide replication restart. J Biol Chem, 2013. 288(42): p. 30139–50.

18. Wang, F., et al., Human CST has independent functions during telomere duplex replication and C-strand fill-in. Cell Rep, 2012. 2(5): p. 1096–103.

19. Feng, X., et al., CTC1-mediated C-strand fill-in is an essential step in telomere length maintenance. Nucleic Acids Res, 2017. 45(8): p. 4281–4293.

20. Boccardi, V., et al., Stn1 is critical for telomere maintenance and long-term viability of somatic human cells. Aging Cell, 2015. 14(3): p. 372–81.

21. Wang, F., J. Stewart, and C.M. Price, Human CST abundance determines recovery from diverse forms of DNA damage and replication stress. Cell Cycle, 2014. 13(22): p. 3488–98.

22. Chastain, M., et al., Human CST Facilitates Genome-wide RAD51 Recruitment to GC-Rich Repetitive Sequences in Response to Replication Stress. Cell Rep, 2016. 16(5): p. 1300–1314.

23. Sun, J., et al., Stn1-Ten1 is an Rpa2-Rpa3-like complex at telomeres. Genes Dev, 2009. 23(24): p. 2900–14.

24. Bhattacharjee, A., et al., STN1 OB Fold Mutation Alters DNA Binding and Affects Selective Aspects of CST Function. PLoS Genet, 2016. 12(9): p. e1006342.

25. Chen, R. and M.S. Wold, Replication protein A: single-stranded DNA’s first responder: dynamic DNA-interactions allow replication protein A to direct single-strand DNA intermediates into different pathways for synthesis or repair. Bioessays, 2014. 36(12): p. 1156–61.

26. Bhattacharjee, A., et al., Dynamic DNA binding, junction recognition and G4 melting activity underlie the telomeric and genome-wide roles of human CST. Nucleic Acids Res, 2017. 45(21): p. 12311–12324.

27. Hom, R.A. and D.S. Wuttke, Human CST Prefers G-Rich but Not Necessarily Telomeric Sequences. Biochemistry, 2017. 56(32): p. 4210–4218.

28. Rhodes, D. and H.J. Lipps, G-quadruplexes and their regulatory roles in biology. Nucleic Acids Res, 2015. 43(18): p. 8627–37.

29. Huppert, J.L. and S. Balasubramanian, Prevalence of quadruplexes in the human genome. Nucleic Acids Res, 2005. 33(9): p. 2908–16.

30. Vannier, J.B., et al., RTEL1 dismantles T loops and counteracts telomeric G4-DNA to maintain telomere integrity. Cell, 2012. 149(4): p. 795–806.

31. Li, X.M., et al., Guanine-vacancy-bearing G-quadruplexes responsive to guanine derivatives. Proc Natl Acad Sci U S A, 2015. 112(47): p. 14581–6.

32. Guilbaud, G., et al., Local epigenetic reprogramming induced by G-quadruplex ligands. Nat Chem, 2017. 9(11): p. 1110–1117.

33. Zimmer, J., et al., Targeting BRCA1 and BRCA2 Deficiencies with G-Quadruplex-Interacting Compounds. Mol Cell, 2016. 61(3): p. 449–60.

34. Paeschke, K., et al., Pif1 family helicases suppress genome instability at G-quadruplex motifs. Nature, 2013. 497(7450): p. 458–62.

35. Sarkies, P., et al., Epigenetic instability due to defective replication of structured DNA. Mol Cell, 2010. 40(5): p. 703–13.

36. Ray, S., et al., RPA-mediated unfolding of systematically varying G-quadruplex structures. Biophys J, 2013. 104(10): p. 2235–45.

37. Mergny, J.L. and L. Lacroix, UV Melting of G-Quadruplexes. Curr Protoc Nucleic Acid Chem, 2009. Chapter 17: p. Unit 17 1.

38. Wang, F., et al., Telomere- and telomerase-interacting protein that unfolds telomere G-quadruplex and promotes telomere extension in mammalian cells. Proc Natl Acad Sci U S A, 2012. 109(50): p. 20413–8.

39. Wan, M., et al., OB fold-containing protein 1 (OBFC1), a human homolog of yeast Stn1, associates with TPP1 and is implicated in telomere length regulation. J Biol Chem, 2009. 284(39): p. 26725–31.

40. Batzer, M.A. and P.L. Deininger, Alu repeats and human genomic diversity. Nat Rev Genet, 2002. 3(5): p. 370–9.

41. Casteel, D.E., et al., A DNA polymerase-{alpha}{middle dot}primase cofactor with homology to replication protein A-32 regulates DNA replication in mammalian cells. J Biol Chem, 2009. 284(9): p. 5807–18.

42. Biffi, G., et al., Quantitative visualization of DNA G-quadruplex structures in human cells. Nat Chem, 2013. 5(3): p. 182–6.

43. Fleming, A.M., et al., 8-Oxo-7,8-dihydroguanine in the Context of a Gene Promoter G-Quadruplex Is an On-Off Switch for Transcription. ACS Chem Biol, 2017. 12(9): p. 2417–2426.

44. Crabbe, L., et al., Defective telomere lagging strand synthesis in cells lacking WRN helicase activity. Science, 2004. 306(5703): p. 1951–3.

45. Maestroni, L., S. Matmati, and S. Coulon, Solving the Telomere Replication Problem. Genes (Basel), 2017. 8(2).

46. Zhao, Y., et al., Processive and distributive extension of human telomeres by telomerase under homeostatic and nonequilibrium conditions. Mol Cell, 2011. 42(3): p. 297–307.

47. Gilson, E. and V. Geli, How telomeres are replicated. Nat Rev Mol Cell Biol, 2007. 8(10): p. 825–38.

48. Schiavone, D., et al., Determinants of G quadruplex-induced epigenetic instability in REV1-deficient cells. EMBO J, 2014. 33(21): p. 2507–20.

49. Li, F., et al., The BUB3-BUB1 Complex Promotes Telomere DNA Replication. Mol Cell, 2018. 70(3): p. 395–407 e4.

50. Nera, B., et al., Elevated levels of TRF2 induce telomeric ultrafine anaphase bridges and rapid telomere deletions. Nat Commun, 2015. 6: p. 10132.

51. Sarek, G., et al., TRF2 recruits RTEL1 to telomeres in S phase to promote t-loop unwinding. Mol Cell, 2015. 57(4): p. 622–635.

52. Sfeir, A., et al., Mammalian telomeres resemble fragile sites and require TRF1 for efficient replication. Cell, 2009. 138(1): p. 90–103.

53. d’Alcontres, M.S., et al., TopoIIalpha prevents telomere fragility and formation of ultra thin DNA bridges during mitosis through TRF1-dependent binding to telomeres. Cell Cycle, 2014. 13(9): p. 1463–81.

54. Sirbu, B.M., et al., Identification of proteins at active, stalled, and collapsed replication forks using isolation of proteins on nascent DNA (iPOND) coupled with mass spectrometry. J Biol Chem, 2013. 288(44): p. 31458–67.

55. Trenz, K., et al., ATM and ATR promote Mre11 dependent restart of collapsed replication forks and prevent accumulation of DNA breaks. EMBO J, 2006. 25(8): p. 1764–74.

56. Georgescu, R.E., et al., Replisome mechanics: lagging strand events that influence speed and processivity. Nucleic Acids Res, 2014. 42(10): p. 6497–510.

57. Maizels, N. and L.T. Gray, The G4 genome. PLoS Genet, 2013. 9(4): p. e1003468.

58. Abd El-Moneam, N.M., et al., Protective role of antioxidants capacity of Hyrtios aff. Erectus sponge extract against mixture of persistent organic pollutants (POPs)-induced hepatic toxicity in mice liver: biomarkers and ultrastructural study. Environ Sci Pollut Res Int, 2017. 24(27): p. 22061–22072.

59. Joo, H.N., et al., pH-Dependant fluorescence switching of an i-motif structure incorporating an isomeric azobenzene/pyrene fluorophore. Bioorg Med Chem Lett, 2017. 27(11): p. 2415–2419.

60. Leon-Ortiz, A.M., J. Svendsen, and S.J. Boulton, Metabolism of DNA secondary structures at the eukaryotic replication fork. DNA Repair (Amst), 2014. 19: p. 152–62.

61. Lin, W., et al., Mammalian DNA2 helicase/nuclease cleaves G-quadruplex DNA and is required for telomere integrity. EMBO J, 2013. 32(10): p. 1425–39.

62. Zaug, A.J., E.R. Podell, and T.R. Cech, Human POT1 disrupts telomeric G-quadruplexes allowing telomerase extension in vitro. Proc Natl Acad Sci U S A, 2005. 102(31): p. 10864–9.

63. Salas, T.R., et al., Human replication protein A unfolds telomeric G-quadruplexes. Nucleic Acids Res, 2006. 34(17): p. 4857–65.

64. Marechal, A. and L. Zou, RPA-coated single-stranded DNA as a platform for post-translational modifications in the DNA damage response. Cell Res, 2015. 25(1): p. 9–23.

65. Sauer, M. and K. Paeschke, G-quadruplex unwinding helicases and their function in vivo. Biochem Soc Trans, 2017. 45(5): p. 1173–1182.

66. Zimmermann, M., et al., TRF1 negotiates TTAGGG repeat-associated replication problems by recruiting the BLM helicase and the TPP1/POT1 repressor of ATR signaling. Genes Dev, 2014. 28(22): p. 2477–91.

67. Anderson, B.H., et al., Mutations in CTC1, encoding conserved telomere maintenance component 1, cause Coats plus. Nat Genet, 2012. 44(3): p. 338–42.

68. Simon, A.J., et al., Mutations in STN1 cause Coats plus syndrome and are associated with genomic and telomere defects. J Exp Med, 2016. 213(8): p. 1429–40.

69. Walne, A.J., et al., Mutations in the telomere capping complex in bone marrow failure and related syndromes. Haematologica, 2013. 98(3): p. 334–8.

